# Unidirectional response to bidirectional selection on body size. I. Phenotypic, life history and endocrine response

**DOI:** 10.1101/498683

**Authors:** Clémentine Renneville, Alexis Millot, Simon Agostini, David Carmignac, Gersende Maugars, Sylvie Dufour, Arnaud Le Rouzic, Eric Edeline

## Abstract

Anthropogenic perturbations such as harvesting often select against a large body size and are predicted to induce rapid evolution towards smaller body sizes and earlier maturation. However, body-size evolvability and, hence, adaptability to anthropogenic perturbations remain seldom evaluated in wild populations. Here, we use a laboratory experiment over 6 generations to measure the ability of wild-caught medaka fish (*Oryzias latipes*) to evolve in response to bidirectional size-dependent selection mimicking opposite harvest regimes. Specifically, we imposed selection against a small body size (Large line), against a large body size (Small line) or random selection (Control line), and measured correlated responses across multiple phenotypic, life-history and endocrine traits. As expected, the Large line evolved faster somatic growth and delayed maturation, but also evolved smaller body sizes at hatch, with no change in average levels of pituitary gene expressions of luteinizing, follicle-stimulating or growth (GH) hormones. In contrast, the Small medaka line was unable to evolve smaller body sizes or earlier maturation, but evolved smaller body sizes at hatch and showed marginally-significant signs of increased reproductive investment, including larger egg sizes and elevated pituitary GH production. Natural selection on medaka body size was too weak to significantly hinder the effect of artificial selection, indicating that the asymmetric body-size response to size-dependent selection reflected an asymmetry in body-size evolvability. Our results show that trait evolvability may be contingent upon the direction of selection, and that a detailed knowledge of trait evolutionary potential is needed to forecast population response to anthropogenic change.

## INTRODUCTION

Human activities often converge towards selecting against large-bodied individuals in animal populations, mainly through harvesting, habitat fragmentation and climate warming (Edeline, 2016). In this context, the dynamics of wild populations may critically rely on their capacity to evolve in response to size-dependent selection.

Whether and how wild populations can respond to anthropogenic size-dependent selection has been mostly explored in the context of fisheries, which are often highly size-selective (Lagler, 1968; Law, 2000; Carlson *et al.,* 2007; Kuparinen *et al.,* 2009). Harvesting large-bodied individuals is predicted to induce adaptive evolution towards earlier maturation through reduced life expectancy and, at the same time, towards slower somatic growth through selection against a large body size at a given age (Heino *et al.,* 2015). Paradoxically, however, selection for an earlier maturation may also result in evolution of faster somatic growth, which allows for an earlier maturation (Dunlop *et al.,* 2009; Eikeset *et al.,* 2016; Diaz Pauli *et al.,* 2017). This result highlights the importance of considering trait correlations and multivariate phenotypes in evolutionary biology.

In the wild, fishing has been associated with phenotypic changes towards earlier maturation at a smaller body size and/or towards slower growth rates (see reviews by Trippel, 1995; Law, 2000; Kuparinen & Merilä, 2007; Fenberg & Roy, 2008; Heino *et al.,* 2015). Yet, cases of stocks with no phenotypic response to fishing are also reported (Devine & Heino, 2011; Silva *et al.,* 2013; Marty *et al.,* 2014), suggesting that harvested populations might not always be able to respond to harvest-induced selection. Studies based on data from the wild, however, are often criticized for problems in measuring actual selection pressures (but see Carlson *et al.,* 2007; Edeline *et al.,* 2007; Kendall *et al.,* 2009), in disentangling the effects on mean trait values of size-selective mortality *vs*. evolutionary changes (Hairston *et al.,* 2005), or in controlling for the confounding effects of phenotypic plasticity (Heino *et al.,* 2002). Hence, there is still debate as to whether changes (or absence thereof) towards earlier maturation and slower somatic growth in exploited populations are genetic (Borrell, 2013), or are occurring rapidly enough to influence population dynamics and thus probability of population persistence (Diaz Pauli & Heino, 2014). Experimental harvesting experiments in the laboratory are potentially free of such problems because they make it possible to accurately target the traits under selection, to fully control the pattern and intensity of artificial selection, as well as to standardize environmental variation so that the effects of phenotypic plasticity are alleviated.

Size-selective experiments have been performed on model organisms such as *Drosophila melanogaster* (e.g., Partridge *et al.,* 1999), chicken *Gallus gallus* (Dunnington *et al.,* 2013) or mice *Mus musculus* (e.g., Macarthur, 1949). Often, selection is bidirectional, i.e., is performed at random (Control line), against a small body size (Large line) and against a large body size (Small line, mimicking the effects of harvesting). Results from these experiments show that body-size response to selection may sometimes be asymmetric, with either the Large or Small lines showing slower, or sometimes no or halted response to selection (Falconer & Mackay, 1996 and references therein; Dunnington *et al.,* 2013; Lynch & Walsh, 2018 and references therein). Additionally, selection on body size may be associated with changes in other traits. For instance, selection for increased thorax length in *D. melanogaster* was associated with an increase in larval development time and no change in somatic growth rate, while selection for reduced thorax length was associated with reduced growth rate but no change in duration of larval development (Partridge *et al.,* 1999). Similarly, experiments specifically designed to simulate harvesting on wild populations of model or non-model organisms have shown that size-at-age or size at maturity in populations subject to small-vs. large-sized harvesting may (Edley & Law, 1988; Conover & Munch, 2002; Amaral & Johnston, 2012; Cameron *et al.,* 2013; van Wijk *et al.,* 2013), or may not (Uusi-Heikkilä *et al.,* 2015) evolve in the direction imposed by selection (see the Discussion for a more detailed treatment of these harvest-simulating experiments). Hence, so far our knowledge of whether and how exploited populations can respond to size-selective harvesting remains limited.

To contribute filling this gap in our knowledge, we examined the ability of a wild population of medaka fish (*Oryzias latipes*) to respond to bidirectional size-dependent harvesting in the laboratory. Specifically, we selected medaka randomly (Control line), against a large body size (Small line), and against a small body size (Large line) during 2.5 years (30 months, 6 medaka generations), measuring at each generation a total of 14 phenotypic, life-history and neuroendocrine traits (Table 1).

**Table 1.**
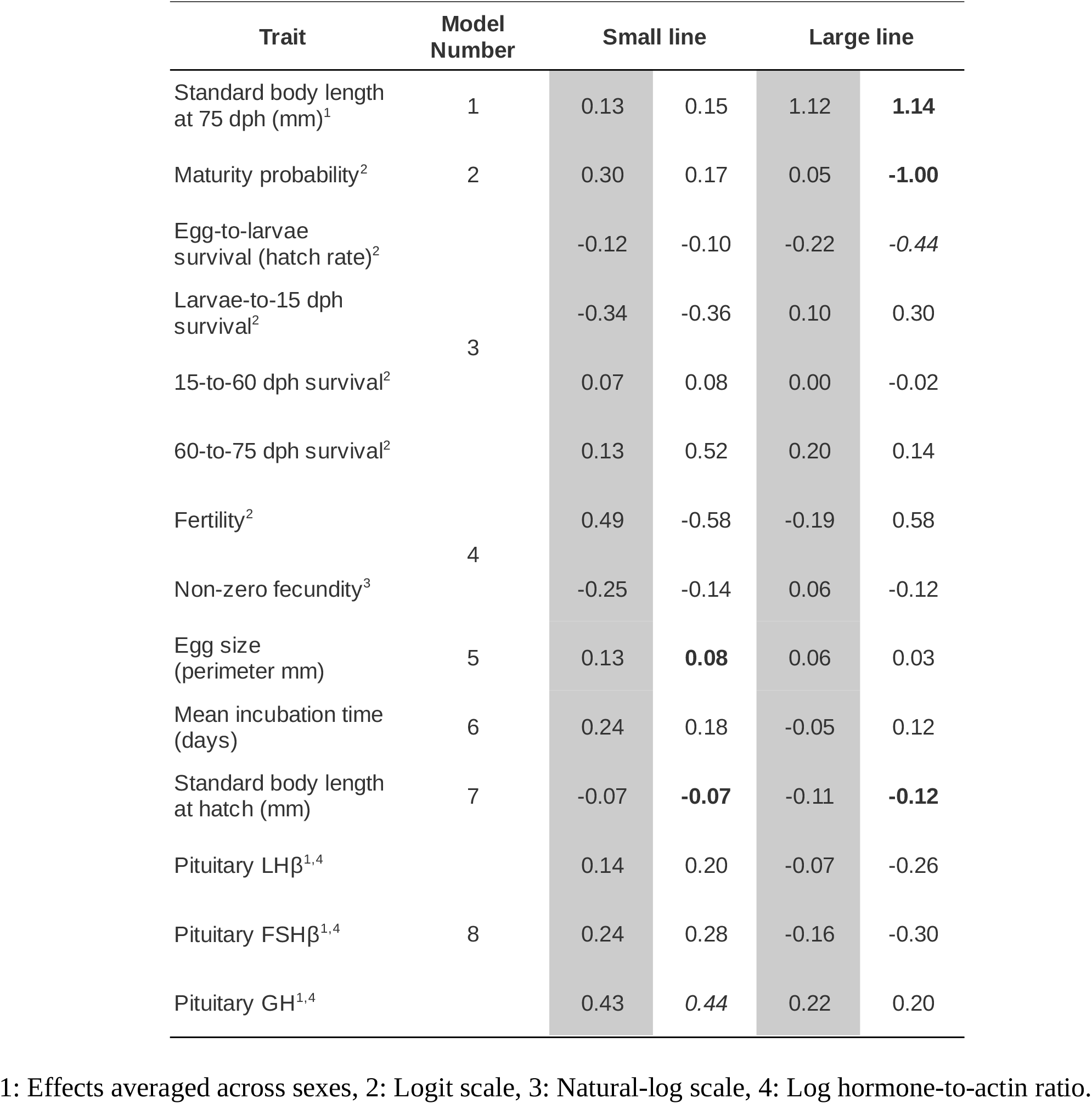
Effect sizes of bidirectional selection on body size on phenotypic, life-history and neuroendocrine traits in medaka. Effects sizes were computed as *μ_S_–μ_C_*, where *μ_S_* and *μ_C_* are mean trait values in Small/Large and Control lines, respectively. Shaded columns show “raw” effect sizes computed from a simple line contrast. Non-shaded columns show effects sizes corrected for the effect of covariates in models 1-8. We tested for significance of the 28 corrected effect sizes by applying a Bonferonni correction in which the significance cut off was α = 0.05/28 = 0.002. Statistically significant values are highlighted in bold. Marginally significant values (α < 0.05) are italicized.

We made three specific predictions for medaka response to size-dependent selection: (1) compared to the Control line, medaka from the Small line should evolve slower somatic growth rates. We predicted an opposite pattern in the Large medaka line. (2) Selection on body size has often been shown to induce correlated responses of reproductive traits and larval viability (e.g., Walsh *et al.,* 2006). Therefore, we predicted that evolution of somatic growth in the Small medaka line should be paralleled by evolution towards increased reproductive investment, which may result in earlier maturation and/or higher fecundity at a given body size and/or larger egg sizes (Roff, 1992), and/or towards reduced size at hatch and larval survival (Walsh *et al.,* 2006). We predicted an opposite response in the Large medaka line. (3) The neuroendocrine control of vertebrate body growth and reproduction involves production of the growth (GH), luteinizing (LH) and follicle-stimulating (FSH) hormones in the pituitary (Rousseau & Dufour, 2007; Zohar *et al.,* 2010). Hence, compared to Control line we predicted altered GH, LH, and FSH expression levels in the pituitary, with potentially opposite alteration patterns in the Small and Large medaka lines. Our results validate prediction (1), but in the Large medaka line only, because the Small line did not show *any* body-size response to selection. Prediction (2) was validated in the Large line, but only partially in the Small line that did not mature earlier but showed signs of increased reproductive investment. Finally, prediction (3) was mainly not supported since only the pituitary expression GH showed a marginally-significant response to size-dependent selection.

## MATERIALS AND METHODS

### Fish origin and maintenance

Our start medaka population descended from 100 wild-caught individuals sampled from a single population in Kiyosu (Toyohashi, Aichi Prefecture, Japan) in June 2011. The genome of the Kiyosu population is free of any significant structure and shows a high degree of polymorphism, indicating no recent population bottleneck (Spivakov *et al.,* 2014). These 100 breeders were maintained in five 20 L aquariums and eggs were collected daily from July to September 2011. Hatched larvae were stocked in six 10 m^3^ outdoor ponds.

In 2013, around 60 adult fish were transferred from outdoor ponds to the laboratory where all the 9 subsequent generations (dubbed F_-1_ to F_7_) were maintained under constant environmental conditions (common garden): 3 L aquariums connected to a continuous flow-through system ensuring good water quality, cycle of 14h of light - 10h of darkness, temperature maintained between 26 and 27.5°C. Fish were fed *ad libitum* with a mixed diet of dry food (Marin Start, Le Gouessant Aquaculture) delivered 4 times per day using automatic microfeeders (Eheim 3581), and live food (*Artemia salina* nauplii and/or *Turbatrix aceti*) manually delivered once a day, 5 days per week. These light, temperature and food conditions provide optimal growth and maturation conditions to medaka (Kinoshita *et al.*, 2009).

### Breeding design, pedigree and fish numbers

Prior to starting selection, we bred medaka during two generations in the laboratory to alleviate maternal and grand maternal effects (a diagram of the experimental design is provided in Appendix 2). Fish initially transferred from outdoor ponds to the laboratory were allowed to mate randomly in groups of 3-6 fish per aquarium to produce the F_-1_ generation. In F_-1_ and F_0_, we randomly mated 54 (F_-1_) and 56 (F_0_) pairs, respectively (Appendix 2), to break any genetic structure or linkage disequilibrium that could remain from possible assortative mating in the wild population (Lynch & Walsh, 2018). Each generation, eggs from each breeding pair were pooled for incubation and larvae from the same clutch were transferred to the same growth aquarium so as to form sibling families. This way, we were able to keep track of individual pedigrees and to estimate individual inbreeding rate as 2k-1, where k is one’s kinship coefficient with oneself (as calculated from the pedigree data using the kinship2 R package, Sinnwell *et al.,* 2014).

Offspring from multiple breeding pairs were never mixed in the same aquarium (not to break the pedigree), and the aquarium and the sibling family effects were confounded. Occasionally, a breeding pair produced many progeny that were spread across two different aquariums (118 breeding pairs, out of 375 pairs in total, produced two aquariums of progeny). Aquariums were randomly spread across two different racks such that the selected lines shared the same environmental conditions. Larvae were initially introduced in their aquariums at a controlled density of 19.6 ± 1.6, 19.2 ± 1.9, 19.8 ± 1.0 (mean ± SD) larvae per aquarium in the Control, Small and Large lines, respectively. Highest densities were suppressed at two weeks post-hatch to reach 17.0 ± 2.3, 16.1 ± 2.1, 17.7 ± 2.0 individuals per aquarium. Densities were not manipulated at later ages. At 76.7 ± 4.4 SD days post-hatch (hereafter 75 dph for short), densities were 15.0 ± 2.4, 14.2 ± 2.1, 15.6 ± 2.4 in the Control, Small and Large lines, respectively.

### Selection procedure

We proceeded with selection on the F_1_ to F_7_ generations from February 2014 to August 2016 (30 months, Appendix 2). A size-dependent selection differential was applied both on families at 60 dph and on mature individuals at 75 dph, an age at which 86% of the fish were mature on average (for dynamics of maturity in each line, see Le Rouzic et al. Under Review).

At 60 dph, we discarded families of less than 10 individuals to avoid confounding density effects on phenotypes. This procedure generated significant selection for a higher fecundity (overdispersed Bernoulli GLM, discarded ~ fecundity, p-value < 0.01) and for higher survival rate from egg to age 15 dph (p-value < 0.005), but not for a larger or smaller body length (p-value = 0.296). Among the remaining families, we kept 10 families at random (Control line) or that had the smallest (Small line) or largest (Large line) average standard body length.

At 75 dph, we individually-selected breeders among mature fish based on their individual standard body length and precluded brother-sister mating. Specifically, we kept in each family 4 mature fish (2 males and 2 females) that were paired with breeders from other families to form the subsequent generation (20 breeding pairs/line/generation, Appendix 2). We formed breeding pairs so as to minimize inbreeding using a computer resampling procedure (selection of the pairing pattern minimizing the median inbreeding coefficient). Assuming no inbreeding in F_1_, mean inbreeding rate was 9.6% (± 1.9 SD) by F_7_. This corresponds to an average effective population size (“inbreeding effective numbers” *sensu* Crow & Kimura 1970 of N_e_ = 30.2).

Each generation, selection was performed on 636 fish on average (212 fish/line), and the selection procedure resulted in keeping on average 12% of individuals per line (number of breeders / total number of fish before selection at 75 dph). We calculated the resultant selection differentials as the difference in maturity probability (i.e., proportion of mature fish) and standard body length after and before selection. Selection differentials across generations F_1_ to F_6_ for maturity proportion and standatd body length were +0.13 (0.12 SD) and +0.68 mm (0.18 mm SD) in the Control line, +0.10 (0.08 SD) and −1.06 mm (0.55 mm SD) in the Small line, and +0.13 (0.08 SD) and +2.05 mm (0.55 mm SD) in the Large Line, respectively.

### Phenotyping

Eggs from each breeding pair were collected during a period corresponding to mother’s 88 to 92 dph. Eggs were counted and photographed, and ImageJ was then used to measure their individual egg perimeters (9795 eggs measured from F_1_ to F_7_). Hatched larvae were collected during a 5-day time window so as to synchronize hatching dates as much as possible. Birthdate was the median hatching date of each sibling family, and all siblings were thus assigned the same age.

At 0 (hatching), 15, 60 and 75 dph each single individual was photographed, and then ImageJ was used to measure standard body length (from the tip of the snout to the base of the caudal fin, 16808 individual measurements from F_1_ to F_7_). Additionally, each individual at each phenotyping was sexed as immature (I), female (F) or male (M) according to their secondary sexual characters (Yamamoto, 1975), which was a non-destructive proxy for the onset of maturity. All fish manipulations were performed after anaesthesia in tricaine methane sulfonate (MS222), except at 0 and 15 dph when larvae and juveniles were manipulated with a Pasteur pipette and photographed in a droplet.

### Pituitary expression of candidate genes

An enzyme-linked immunosorbent assay (ELISA) is not available for medaka GH and ELISAs, in addition of being much less sensitive than reverse transcription quantitative real-time polymerase chain reaction (RT-qPCR), require plasma volumes that are too large to allow individual measurements in medaka. Hence, as a first approach to uncovering the molecular regulation of adaptive life-history evolution in medaka, we used RT-qPCR to measure mRNA levels of candidate genes in individual pituitaries. Specifically, we measured pituitary mRNA levels of β-subunits of gonadotropin hormones (LHβ and FSHβ) and GH. F_0_ preliminary data indicated that the onset of secondary sexual characteristics occurred roughly between 40 and 60 dph, and we chose to dissect fish at 40 dph so as to sample fish at the initiation of puberty. In each generation from F_1_ to F_7_, 10 to 15 fish per line (233 fish in total) were phenotyped as described above, sacrificed and dissected under a binocular microscope for the pituitary which was immediately immersed in 250 μL Trizol (Ambion) and stored at −20°C.

After sample homogenization by agitation (15 sec vortexing), total RNA were extracted according to the manufacturer’s indications, suspended in 10 μL RNAse-free water, and treated with DNAse I (Dnase I recombinant RNAse-free, Roche Diagnostics). Then cDNA were produced from 5 μL of total RNA using RT Superscript III (RT Superscript III First Strand cDNA Synthesis Kit; Invitrogen, Life Technologies) and random hexamer primers (50 ng; Invitrogen, Life Technologies), at 50°C for 60 min after an initial step of 25°C for 10 min. Medaka specific primer sets for FSH were designed with primer3 software (Koressaar & Remm, 2007) on two successive exons, or on exon junctions. Gene specific primer sets for LHβ, FSHβ, GH and actin-β (used as reference gene to correct for technical noise), were previously designed (see Appendix 1). Efficiency and amplification specificity were checked for each primer set. The sets with the highest efficiency were chosen for the following quantification experiment.

Messengers RNAs were assayed using Light Cycler system (LightCycler® device; Roche Diagnostics) with the LightCycler FastStart Master plus SYBR Green I kit (Roche Diagnostics) as recommended by the manufacturer, from 4 μL of diluted 1:10 cDNA samples and the specific primers concentrated at 500 nM (Eurofins). The PCR conditions were 95°C for 10 min followed by 50 cycles at 95°C for 5 sec, 60°C for 10 sec and 72°C for 5 sec.

Expression levels of mRNA for LHβ, FSHβ, GH and actin-β in each individual fish were measured in duplicate using the “relative quantification” method (Applied Biosystems User Bulletin #2). Briefly, the standard relationships between fluorescence and gene-specific sample RNA concentrations were constructed using a bulk RNA pool, hereafter dubbed “calibrator”. The Lightcycler software estimated the number C_q_ of quantification cycles needed to reach the inflection point (second derivative equal to 0) of fluorescence amplification for a series of 7 calibrator volumes. From this, the software estimated the intercept and slope parameters for the gene-specific, linear relationship between log10 calibrator volume and C_q_. These linear relationships were then used to predict sample-specific mRNA expressions in log10 calibrator volume units (i.e., “arbitrary” units) for LHβ, FSHβ, GH and actin-β from their sample-specific C_q_. At each PCR run, a known amount of the calibrator plus a blank (water) were measured for C_q_ so as to adjust for possible inter-run noise. Following standard practises, we used as input data relative pituitary gene expression, calculated as the natural logarithm of the ratio between mRNA expression for the interest gene and mRNA expression for actin-β (see model 5 below). In particular, this approach corrects for the effects of variability in pituitary size.

### Data analyses

The aim of our statistical analyses was to estimate and test for an overall effect of the selected lines on traits, pooling data from generations F_3_ to F_7_ and treating generation as a random effect. An archive containing datasets and scripts to reproduce analyses is provided as a Supplementary material.

#### Standard body length at 75 dph

We modelled response to selection as the line effect on standard body length at 75 dph (*Sdl* 75) of each individual *i* :

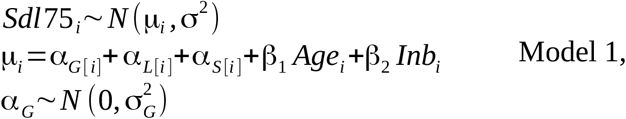

where *N* is the normal distribution, subscripts *G* [*i*], *L* [*i*] and *S* [*i*] denote effects of the generation (F_3_ to F_7_) treated as a normally-distributed random effect, selected line (Small, Large and Control) and sex (I, M or F), respectively. *Age* is age in dph coded as a continuous variable, *Inb* is individual inbreeding coefficient. Finally, σ^2^ is residual variance and 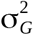 is the variance of the normally-distributed generation effect.

#### Probabilistic maturation reaction norms

We visualized the effect of anthropogenic selection on the maturation process using probabilistic maturation reaction norms (PMRNs). This approach was developed to account for the plastic effects of juvenile somatic growth rate on the maturation process, such that a shift in the maturation reaction norm may be interpreted as an evolutionary shift in maturation (Heino *et al.*, 2002; Heino & Dieckmann, 2008). PMRNs classically account for the effects of age and body length on maturation, but they may also be “higher dimensional” to account for the effects of body mass or individual somatic growth rate (e.g., Morita & Fukuwaka 2006). Here, however, we neither weighed individual medaka nor followed individual growth trajectories. Therefore, we used classical age- and length-dependent PMRNs, which have been demonstrated to be as efficient as higher-dimensional PMRNs to detect evolutionary trends (Dieckmann & Heino, 2007).

For each medaka line, we computed age- and length-dependent PMRNs, defined as the age- and length-dependent 50% probability for an immature medaka to initiate maturity (as informed by the onset of secondary sexual characteristics), using the methods of Barot et al. (2004) and Van Dooren et al. (2005). The methods consisted first in computing maturity “ogives” as:

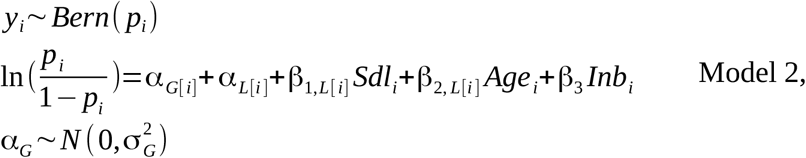

where *y_i_* is the maturity status of an individual fish *i* (0 or 1), *Bern* is the Bernoulli distribution of “success” (maturity) probability *p*, ln is the natural logarithm. Other subscripts or variables are as described above. By letting the effects of both *Sdl* and *Age* on *p* varying for each selected line this model captured potential effects of selection on both the intercept and slope of the PMRN.

Second, we computed the *maturation* probability *m* (*a_τ_,s_τ_*) at each growth increment *τ* as described by Barot et al. (2004):

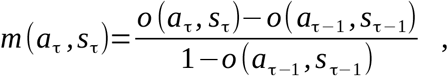

where *o*(*a_τ_,s_τ_*) is age- and length-dependent *maturity* probability at the end of growth increment τ as predicted by model 2. We did so for simulated slow, median and fast growth curves (Van Dooren *et al.*, 2005; Harney *et al.*, 2013).

Finally, we computed line-specific PMRNs as the age and length combination (*a_t_,s_t_*) at witch maturation probability reached 50%, i.e., as the age and length combination that satisfied the following condition (Van Dooren *et al.,* 2005; Harney *et al.,* 2013):

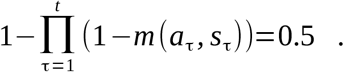

We estimated *m* (*a_τ_, s_τ_*) for 200 growth increments τ equally spread between ages 0 and 87 dph. Full propagation of error distribution for *m* (*a_τ_, s_τ_*) was obtained by iterating the procedure for each Monte Carlo Markov Chain (MCMC) sample of the parameter set in model 2.

#### Survival

We tested for differential mortality among the selected medaka lines using models of the form:

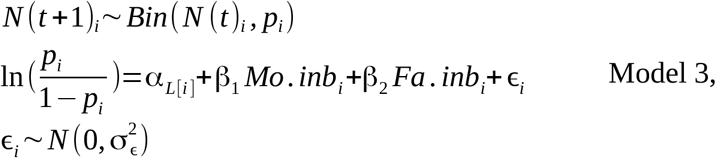

where *N*(*t* + 1)_*i*_ is the number of individuals still alive at time *t* + 1 in sibling family *i*, *Bin* is the Binomial distribution, *Mo. inb* is mother inbreeding coefficient, *Fa. inb* is father inbreeding coefficient. ϵ is an overdispersion effect accounting for the fact that observed variance was larger than canonical variance of the Binomial distribution. We fitted separately four models for *t* to *t* + 1 steps corresponding to the egg-larvae (egg-to-0 dph), larvae-juvenile (0-to-15 dph), juvenile-adult (15-to-60 dph) and adult-adult (60-to-75 dph) transitions.

#### Size-specific fertility and fecundity

Our aim here was to test for possible effects of size-dependent selection on medaka size-specific fecundity. Counts *F* of clutch size per breeding pair *i* were zero-inflated Poisson-distributed and modelled as:

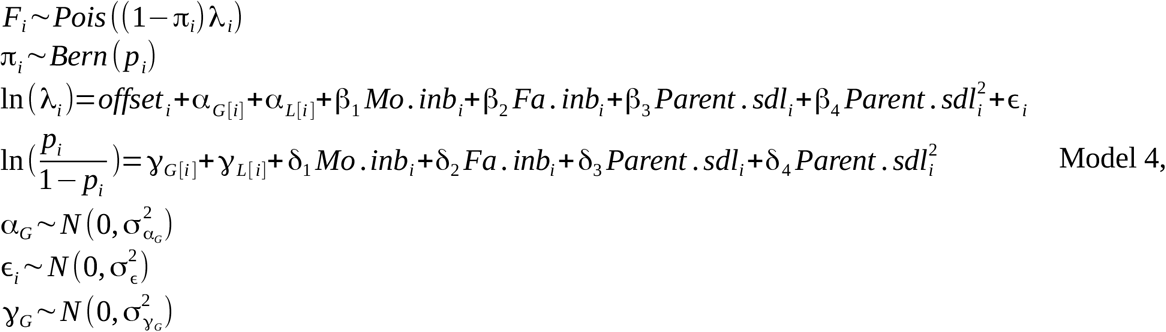

where *Pois* is the Poisson distribution with mean (and variance) equal to the product of probability for a nonzero count 1 – π with nonzero counts λ. This way, we were able to simultaneously test the effects of the predictors both on the probability for a breeding pair to be infertile (*p*) and on the fecundity of a fertile pair (λ).

*offset* is the natural logarithm of number of days during which eggs were collected (varied from 4 to 5 days), *Parent. sdl* is average parent standard body length, and ϵ is an overdispersion effect accounting for the fact that variance overwhelmed the mean in non-zero egg counts. Other subscripts or variables are as described above. In this model, the effect of parent Sdl is accounted for, such that a significant effect of the selected line would indicate that size-dependent selection affects medaka fertility or fecundity beyond direct effects on Sdl.

#### Egg size

Individual egg perimeter *Pm_i_* in mm was modelled as follows:

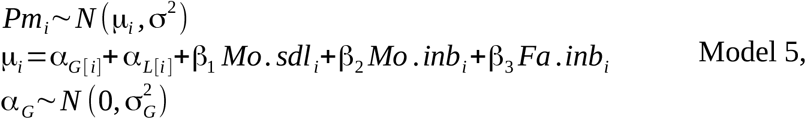

where the variables are as described above in Eqs. 1 and 3.

#### Incubation time

Incubation time *It_i_* for eggs from each breeding pair *i* was computed as the time lapse (days) between mean date of spawning and mean date of hatching for larvae collected from mother’s 95 to 100 dph. We evaluated the effect of selection on *It_i_* in:

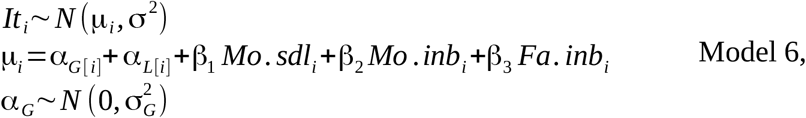

where variables are described above in Eqs. 1 and 3.

#### Larval size at hatch

We modelled individual standard body length at hatch *Sdl* 0 as:

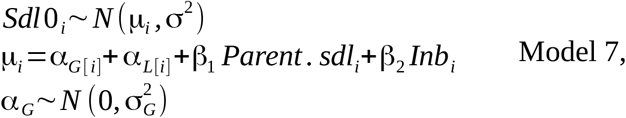

where variables are described above.

#### Hormonal profile

Measurements of LHβ, FSHβ and GH represent a multivariate phenotype recorded on the same individuals. This interdependency of measurements should be accounted for when analysing the effects of selection on traits. Therefore, we modelled the relative pituitary gene expression (see definition above) of individual *i* using a multivariate normal model of the form:

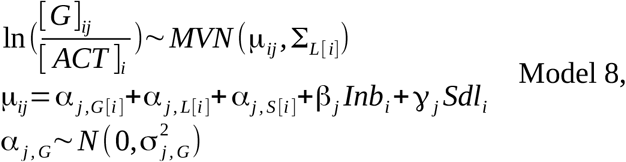

where [*G*]_*ij*_ is the RT-qPCR-measured concentration of the interest gene *j* (LHβ, FSHβ and GH) in the pituitary of individual *i*, [*ACT*]_*i*_ is the RT-qPCR-measured concentration of the reference gene (actin-β) in the pituitary of the same individual *i*, MVN is the multivariate normal distribution, Σ_*L*[*i*]_ is the line-specific variance-covariance matrix of the MVN. We specified an uninformative inverse Wishart prior distribution for Σ_*L*[*i*]_ such as 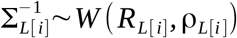 where *W* is the Wishart distribution, *R* is a scale matrix (diagonal matrix of dimension *j*) and ρ = *j* denotes degrees of freedom (Lunn *et al.,* 2012). In practice *R*, which is supplied as data, contains ρ on the diagonal and 0s in non-diagonal entries.

#### Natural selection

Our datasets also allowed us to measure natural selection, which often opposes the effects of artificial selection (e.g., Carlson *et al.,* 2007). In medaka in the laboratory, natural selection may act on the standard body length of the selected parents through affecting their reproductive success or through the survival of their progeny. We visualized these potential effects of natural selection using quadratic regressions of fitness components (namely fecundity, hatch rate, number of progeny reaching age 75 dph, and number of progeny kept as breeders for the next generation) on mean parental body length in linear models (Lande & Arnold, 1983). Specifically, for fecundity, the number of progeny reaching age 75 dph, and number of progeny kept as breeders for the next generation we used a zero-inflated Poisson model similar to Model 4 above, except that fixed effects in linear predictors included only mean parental body length and mean parental body length squared, and that no overdispersion parameter was needed for number of progeny. For modelling hatch rate, we used a binomial model similar to Model 3 above, except that fixed effects included only mean parental body length and mean parental body length squared, and that no overdispersion parameter was needed.

#### MCMC parameter estimation

All models were fitted using MCMC in JAGS (Plummer, 2003) through the jagsUI R package (Kellner, 2019) in R 3.6.3 (R Core Team, 2020). We used weakly informative priors and, for each model, we ran three independent MCMC chains thinned at a period of 5 iterations until parameter convergence was reached, as assessed using the Gelman–Rubin statistic (Gelman & Rubin, 1992).

We tested the significance of effects from posterior parameter distributions using a test equivalent to a two-way *t* test. In these tests, the MCMC P-value was twice the proportion of the posterior for which the sign was opposite to that of the mean posterior value. We further assessed goodness of fit of our models by using a Bayesian P-value (Gelman *et al.,* 1996). Briefly, we computed residuals for the actual data as well as for synthetic data simulated from estimated model parameters (i.e., residuals from fitting the model to ‘‘ideal’’ data). The Bayesian P-value is the proportion of simulations in which ideal residuals are larger than true residuals. If the model fits the data well, the Bayesian P-value is close to 0.5. Bayesian P values for our models ranged from 0.49 to 0.66, indicating excellent model fit. All models were fitted using an “effect” parametrization (Appendix 4), i.e., by setting one level of each factor as a reference levels as is done by default in the R software.

## RESULTS

Effect sizes for response to selection of the 14 measured traits are presented in Table 1, while quantitative statistical results are provided in Appendix 4.

In line with our first prediction, the Large medaka line evolved towards a larger standard body length at 75 dph in both mature (Fig. 1A, Appendix 3) and immature fish (Fig. 1B, Appendix 3). This effect was identical in females, males and immatures at 75 dph (+1.23 mm, MCMC p-value = 0.000, results shown for females only in Model 1 in Appendix 4). However, in contrast with our first prediction, body size in the Small medaka line did *not* respond to selection (Fig. 1, Appendix 3). This lack of response was consistent across females, males and immatures (−0.02 mm, MCMC p-value > 0.800, results shown for females only in Model 1 in Appendix 4). Therefore, medaka presented a unidirectional response to bidirectional size-dependent selection.

**Fig. 1.**
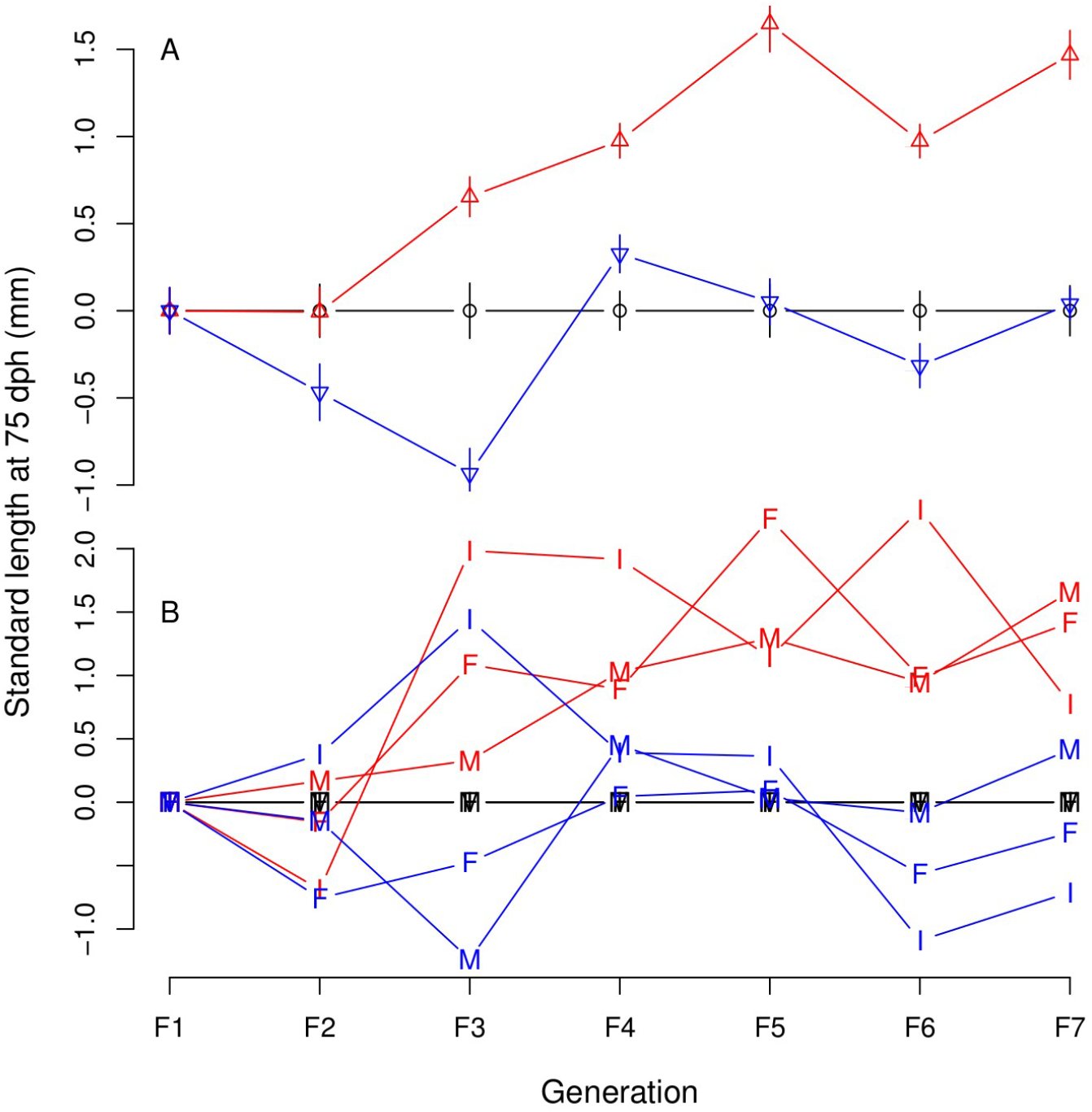
Medaka body-size time series response to bidirectional selection on body size. A: mean standard body length of mature fish (± SE) at 75 dph. Black circles: Control (random size-selected) line; Blue bottom-pointing triangles: Small line; Red, top-pointing triangles: Large line. B: same as A but separately for immature (I), male (M) and female (F) fish and without error bars. Data were centred on the mean of the control line (for raw data, see Appendix 3).

Our second prediction was that evolution of body-size should be paralleled by evolution of correlated traits, and in particular of age and size at maturation, size-specific fecundity, egg sizes, size at hatch and larval survival. Only maturity probability at 75 dph responded as expected, and more sharply so in the Large than in the Small line (Table 1). Specifically, maturity probability at an average age and body length decreased significantly in the Large medaka line only (Model 2 in Appendix 4). This change was associated with an upward shift the probabilistic maturation reaction norm (PMRN) for the Large medaka line compared to the PMRN for the Control line (Fig. 2).

**Fig. 2.**
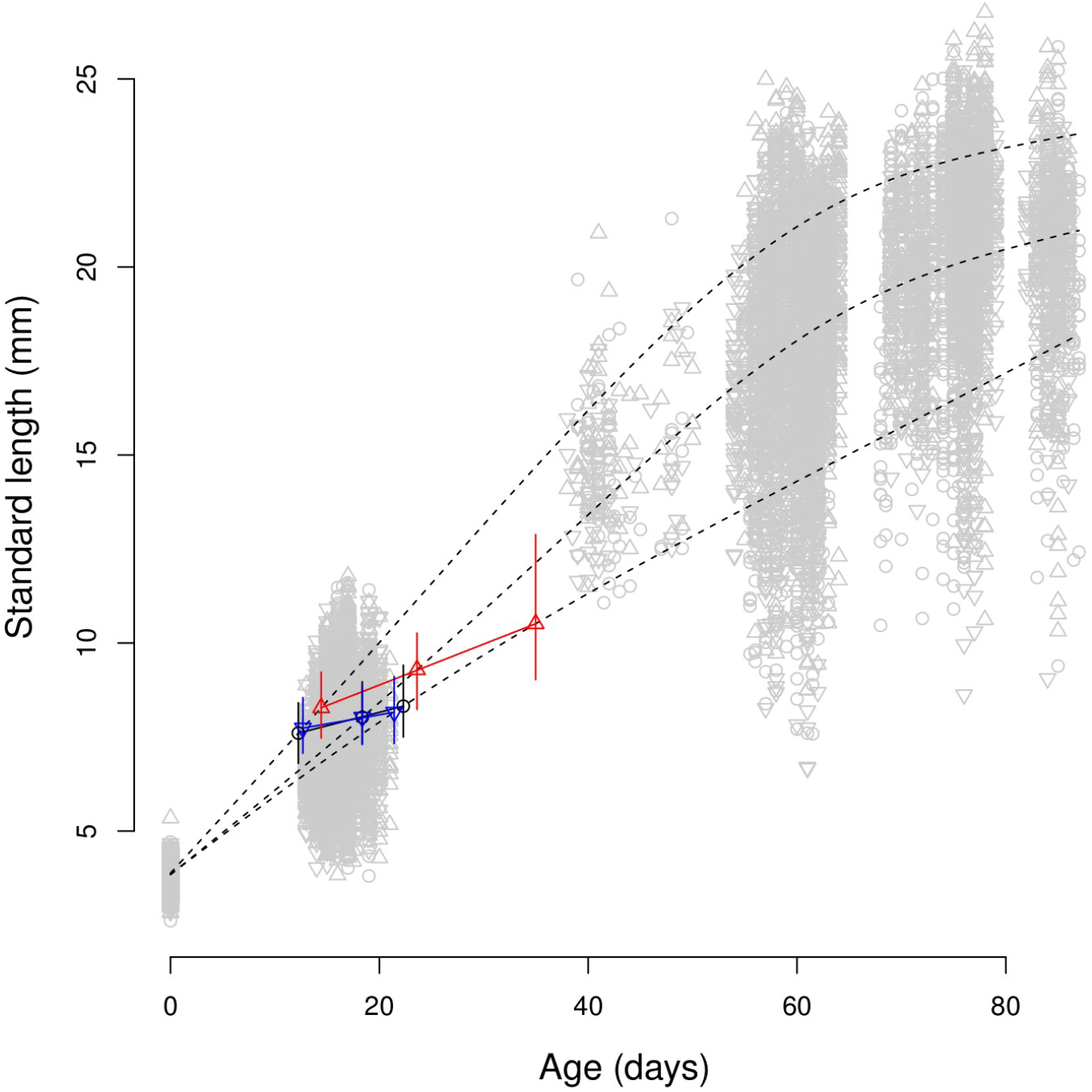
Medaka probabilistic maturation reaction norm (PMRN) response to bidirectional selection on body size. Light grey dots are raw data. Black dotted curves represent simulated slow, medium and fast growth trajectories. Coloured solid lines and dots represent 50% PMRNs and their intersection with the simulated growth curves, respectively. Black circles: Control (random size-selected) line; Blue bottom-pointing triangles: Small line; Red, top-pointing triangles: Large line. Error bars around the coloured dots represent 95% MCMC credible intervals.

In the Small medaka line, maturity probability at an average age and body length did not respond to selection (Model 2 in Appendix 4) and, accordingly, PMRNs for the Small and Control lines largely overlapped (Fig. 2). Noticeably, however, there were some signs of an increased reproductive investment in the Small medaka line: the length-corrected maturity probability decreased less fast with an increasing age than in the Control line (Model 2 in Appendix 4), and egg sizes increased (Table 1, Model 5 in Appendix 4, see also results on GH below).

In contrast with our second prediction, we found that body length at hatch was significantly decreased in *both* the Large and Small medaka lines, as compared to the Control line (Table 1, Model 7 in Appendix 4). This result suggests that larvae might have had larger yolk sacs in these two lines, owing to their similar and larger eggs sizes, respectively. We did not photograph yolk sacs and can not test this hypothesis. Noticeably, body length at hatch was also the only of the 14 monitored traits that was significantly influenced by inbreeding, more inbred individuals having a larger size at hatch (Table 1, Model 7 in and S2). Hatch rate marginally decreased in the Large line compared to the Control line, but we found no effect of selection on survival at later development stages (Table 1, Model 3 in Appendix 4).

Our third prediction was that evolution of body size and maturation should be associated with changes in pituitary production of the growth hormone (GH), and of the β subunits of luteinizing (LH) and follicle-stimulating (FSH) hormones. Mean pituitary expression levels of GH marginally increased in males (but not females) in the Small (but not Large) medaka line compared to the Control line (Fig. 3, Table 1, Model 8 in Appendix 4). There was a trend towards mean pituitary expression levels of LH and FSH to increase in the Small line, and to decrease in the Large line (Fig. 3). However, these trends were not statistically significant (Table 1, Model 8 in Appendix 4), highlighting a probable lack of statistical power. Interestingly, residual pituitary gene expressions for the three hormones did not trade off, but were instead highly positively correlated (Model 8 in Appendix 4). Finally, the positive residual correlation between LH and GH significantly increased in the Large line compared to the Control line (Appendix 5).

**Fig. 3.**
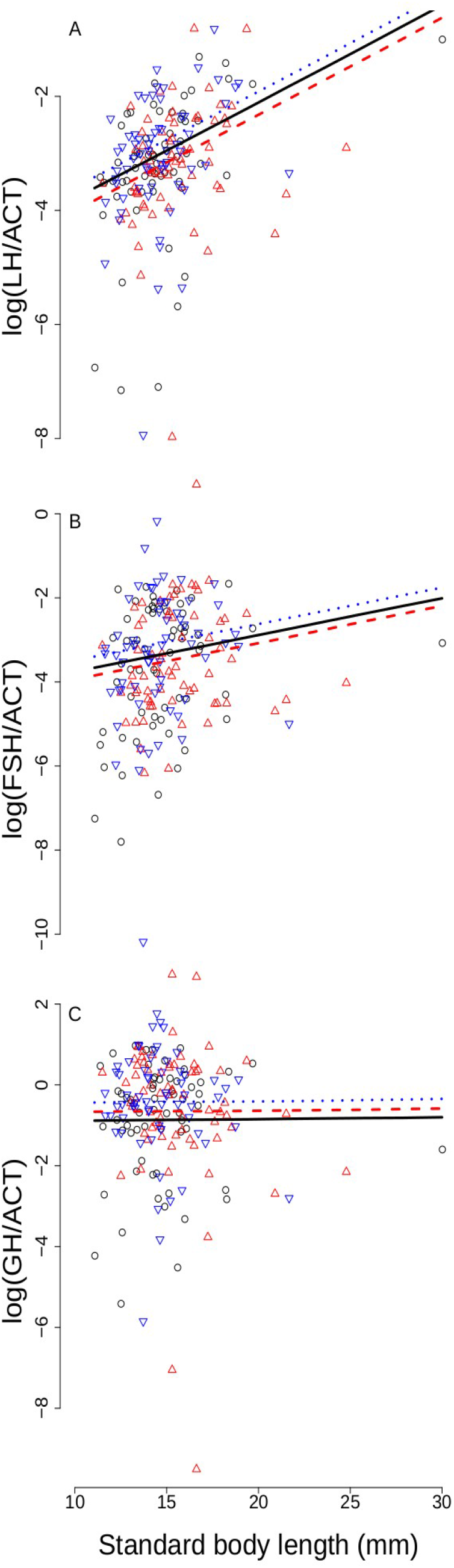
Medaka endocrine response to bidirectional selection on body size. Pituitary mRNA levels for A: the luteinizing hormone (LH, β subunit), B: the follicle-stimulating (FSH, β subunit) and C: growth hormone (GH) were standardized by actin β (ACT) levels and log-transformed. Dots represent raw data. Black circles: Control (random size-selected) line; Blue bottom-pointing triangles: Small line; Red, top-pointing triangles: Large line. Lines represent mean MCMC model predictions. Black solid lines: Control medaka line; Blue dotted lines: Small medaka line; Red dashed lines: Large medaka line. For clarity, only model predictions for males are represented.

We detected significant natural selection on medaka body length during our experiment. Specifically, a longer mean parental body length was associated with increased fecundity (Fig. 4A, effect non statistically significant when inbreeding was also included in Model 4 in Appendix 4), but with a decreased egg hatch rate (Fig. 4B, Table 1, Model 3 in Appendix 4). Despite normalization on higher densities at 15 dph, longer-bodied medaka parents still had an increased number of progeny reaching 75 dph (Fig. 4C) and, despite controlled pairing at 75 dph, stabilizing natural selection on parental body length remained present in terms of number of progeny being selected as breeders for the next generation (Fig. 4D). Therefore, natural selection opposed the effects of artificial selection on medaka body size during our experiment.

**Fig. 4.**
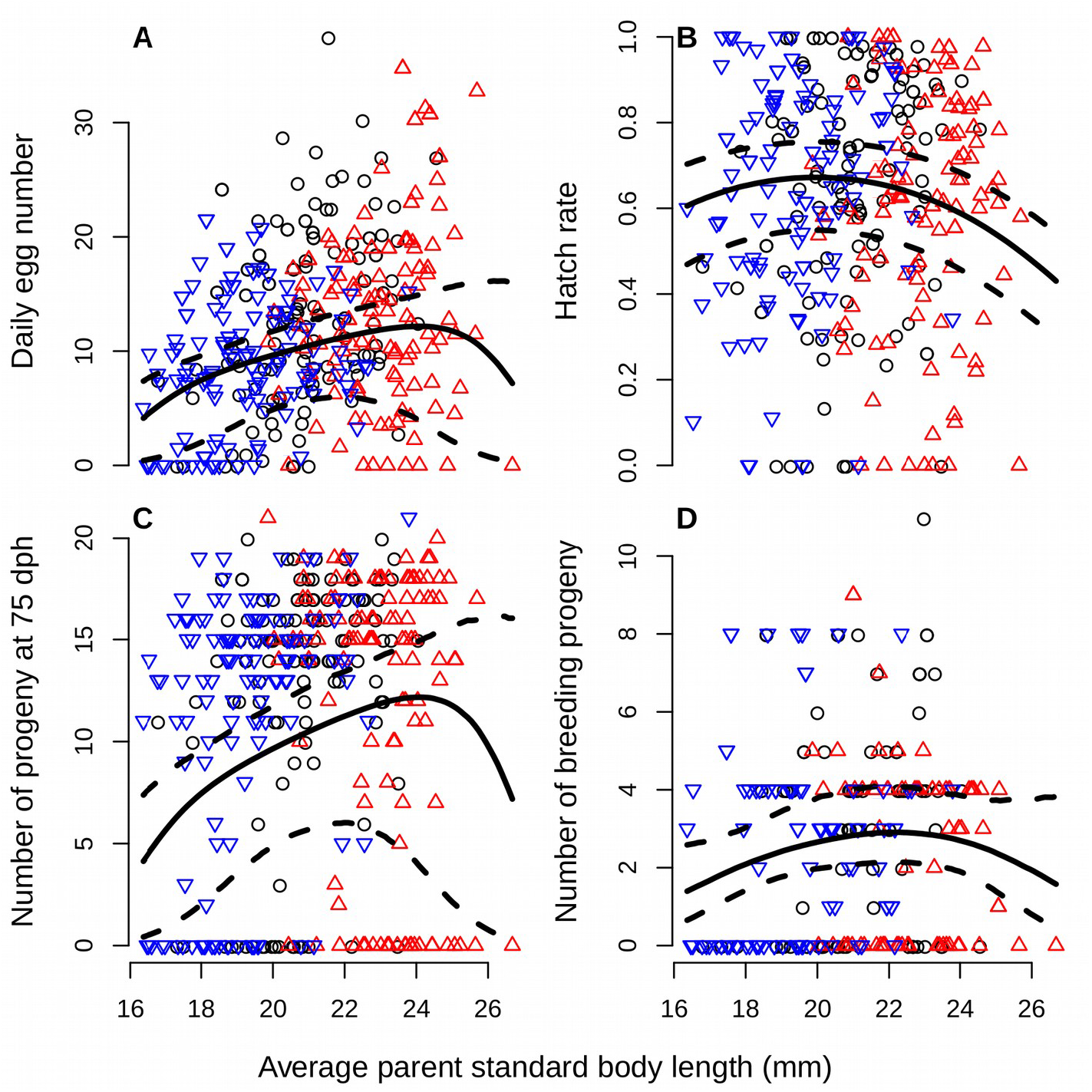
Natural selection on medaka body size in the laboratory. Fitness components are regressed against average standard body length of the parental pair. A: Daily fecundity. B: Hatch rate of the aggregated clutches. C: Number of progeny reaching an age of 75 days-post-hatch. D: Same as C but after the progeny was selected as breeder to produce the next generation. Solid lines show mean MCMC predicted values and dashed lines 95% credible intervals. Dots represent raw data. Black circles: Control (random size-selected) line; Blue bottom-pointing triangles: Small line; Red, toppointing triangles: Large line.

## DISCUSSION

We measured in the laboratory the realized evolvability of body size in response to size-dependent selection in wild-caught medaka fish. We show that medaka responded to selection for a large body size, but not to selection for a small body size. Before discussing this unexpected result, we start with a mini review of previous harvest-simulating experiments and how their results and designs compare to ours.

### Laboratory harvesting experiments

Size-selection experiments are a classic in evolutionary biology, and have been conducted multiple times on model organisms such as mice (e.g., Macarthur, 1949; Falconer, 1973), chicken (Dunnington *et al.,* 2013) or drosophila (e.g., Hillesheim & Stearns, 1991; Partridge *et al.,* 1999). More recently, problems with overexploitation have renewed the interest in size-selective experiments mimicking size-selective harvesting. In a pioneering study, Edley & Law (1988) have applied small *vs*. large harvesting during a 150 day period to six clonal populations of *Daphnia magna.* About 200 individuals were left in each clonal population after each round of harvesting. Populations of clones exposed to small-harvesting (Large lines) evolved rapid somatic growth through small size classes and delayed maturation, while populations of clones exposed to large-harvesting (Small lines) evolved slow growth through small size classes and earlier maturation. Computation of reproductive values showed that evolution resulted in a redistribution of reproductive investment towards size classes that were not harvested.

Conover & Munch (2002) applied small, large or random harvesting at 190 days postfertilization (dpf) during five generations in six experimental populations of the Atlantic silverside *Menidia menidia* maintained in 700L tanks (about 100 breeders/generation/population). The Atlantic silverside is an annual fish, and it was assumed that all individuals were mature at selection such that selection was imposed on body size only. Conover & Munch (2002) found that the mean weight of fish evolved in the expected direction and, by generation F_5_, an average fish aged 190 dpf weighted 3.5 g in the Control lines, 2.5 g in the Small lines, and 4.5 g in the Large lines. These differences were due to differences in somatic growth rate and underlying traits (Walsh *et al.*, 2006).

Amaral & Johnston (2012) applied small, large or random harvesting at 90 dpf on six populations of zebra fish *Danio rerio* maintained in 25 L tanks (24 to 78 breeders/generation/population). After four generations, the selected lines changed in the expected directions with the Small and Large lines evolving mean standard body lengths 2% lower and 10% larger than in the Control line, respectively (actual body length values not presented).

Cameron et al. (2013) exposed soil mites *Sancassania berlesei* to juvenile or adult harvesting during 70 weeks (i.e., harvesting was stage- but not directly size-dependent). There were 6 populations per harvest treatment, plus six unharvested populations (hundreds of individuals per population). In accordance with theoretical predictions (Heino *et al.*, 2015), juvenile harvesting induced evolution towards earlier maturation, while adult harvesting induced evolution towards delayed maturation. Interestingly, the amplitude of harvest-induced evolution was overwhelmed by evolution of delayed maturation in all treatments. This change was interpreted by authors as a response to the captive environment, in which density and competition for resources were increased compared to the natural environment from where mites were initially sampled.

van Wijk et al. (2013) applied small, large or random harvesting in the guppy *Poecilia reticulata* during a 3-generation experiment in five experimental populations maintained in 120L aquariums (125 breeders/generation/population). Male guppy stop growing at maturation, and selection was applied on the body length of mature males only. After 3 generations of selection, body lengths of mature male guppy were on average 21 mm in the Large lines *vs*. 18 mm in the Small lines (19 mm in the Control line). However, the age of males was not standardized, such that it is unclear whether selection acted on male age at maturation, on male somatic growth rate or on both traits simultaneously.

Finally, Uusi-Heikkilä et al. (2015) applied small, large or random harvesting during 5 generations on six experimental populations of zebra fish that were maintained in 320L tanks (120 breeders/generation/population, mating by groups of 2 or 4 fish). Zebra fish were harvested at an age corresponding to 50% of mature fish in the Control line, and breeders were mated 14 days later. Response to selection was contingent upon both the trait considered and upon the direction of selection. Compared to the Control line, the Large line showed no change in juvenile somatic growth rate or asymptotic length but matured at a later age (but not size), while the Small line showed no change in juvenile somatic growth rate but evolved lower asymptotic length and maturation at a smaller size (but not age).

All the above-listed designs, and ours as well, imposed truncation selection on body size, which may or may not accurately reproduce the form of fishing-induced selection depending on the fishing gear. Towed gears and long-lining catch all individuals above a threshold body size, and their effects are thus accurately simulated by truncation selection. In contrast, gillnets or traps selectively target medium-long individuals (Lagler, 1968; Millar & Fryer, 1999; Carlson *et al.,* 2007; Kendall *et al.,* 2009; Kuparinen *et al.,* 2009), and thus generate at the same time disruptive selection and directional selection against a large body size (Carlson *et al.*, 2007; Edeline *et al.*, 2009). Truncation selection does not reproduce the disruptive component of gillnet-induced selection, but it still does capture the directional component. Hence, on the whole truncation selection provides a simple and relatively inclusive selection framework to simulate fishing-induced selection on body size.

Another key feature of all previous laboratory harvesting experiments is that they used a mass-selection design with replication of the selected lines, but no control over effective population sizes, inbreeding rate or natural selection. To avoid these problems, we isolated selected pairs and raised their offspring in individual tanks, keeping track of the pedigrees along the experiment. This made it possible to control for the number of offspring per individuals, to maximize effective population sizes, to limit inbreeding throughout the selection procedure, and to measure natural selection. To our knowledge, this is the first time that such a high level of control is achieved in a size-selection experiment on fish.

However, because the number of individuals included in such an experiment is limited, line replication trades off with increasing effective population size N_e_. A large N_e_ decreases genetic drift, limits the effect of linkage disequilibrium on selection limits, and delays the unavoidable increase in inbreeding (Robertson, 1960; Hill & Robertson, 1966), see e.g. Weber & Diggins (1990) for experimental evidence. In particular, avoiding genetic drift and inbreeding is crucial when studying the evolution of correlated characters (Phillips *et al.*, 2001). Therefore, we chose to derive three large-population lines (25 < N_e_ < 30 in each) rather than replicating small-population treatments. A pedigree-based quantitative genetic analysis suggests that medaka trait dynamics in the Large line were not compatible with random drift, and instead reflected deterministic evolutionary processes (Le Rouzic *et al.*, Under Review).

### Medaka phenotypic and life-history response to bidirectional selection on body size

At the end of our experiment (F_7_), body sizes of mature medaka at 75 days-post-hatch were 20.5 vs. 22.0 mm (7% difference) in the Control vs. Large lines, respectively. This difference is modest, but is in the range of responses to selection observed in other fish harvesting experiments for the Control *vs*. Large lines: 62.3 vs. 76.1 mm (22% difference) in the Atlantic silverside (Conover & Munch, 2002, mean lengths estimated from a mass-length relationship based on data from Duffy *et al.,* 2013), 10% (raw data not available) in zebra fish *Danio rerio* (Amaral & Johnston, 2012), 19.3 vs. 20.8 mm (7.5%) in the guppy *Poecilia reticulata* (van Wijk *et al.,* 2013), and 29.2 vs 29.5 mm for asymptotic length (<1% difference) or 22.6 vs. 22.9 mm for length at maturity (1.2% difference) in zebra fish (Uusi-Heikkilä *et al.*, 2015).

In contrast, medaka body size did not respond to selection in the Small line. Such an unidirectional response to bidirectional selection was not found in previous experiments on Atlantic silverside (Conover & Munch, 2002), zebra fish by Amaral & Johnston (2012) or guppy (van Wijk *et al.,* 2013), but compares with the results obtained on zebra fish by Uusi-Heikkilä et al. (2015), who show that the magnitude of response to size-dependent selection was trait-specific and contingent upon the direction of selection (see above). The qualitative agreement between our results and those of Uusi-Heikkilä et al. (2015) might possibly come from a convergence among our respective selective designs. The selection procedure by Uusi-Heikkilä et al. (2015) involved mating the fish 14 days after that 50% of the population reached maturity, a delay that was possibly not long enough to allow for 100% of the fish to reach maturity, in which case selection was applied *both* on body size and for maturity (similar to our own design). As discussed by Le Rouzic et al. (Under Review), available evidence suggests that response to such bivariate selection on correlated traits is often erratic.

In our experiment, lack of body-size response to selection in the Small medaka line could not be ascribed to an absence of artificial selection, which was strong and consistent, nor due to the counteracting effects of natural selection, which remained weak compared to the strength of artificial selection, nor due to inbreeding which was by F_7_ identical among the random- and large-harvested lines. Instead, the absence of evolution in the Small medaka line suggests that medaka are at a lower evolutionary limit for body size.

### Medaka neuroendocrine response to bidirectional selection on body size

We specifically targeted genes known to play a central role in the regulation of somatic growth and reproduction. In teleosts, growth hormone (GH) is a pleiotropic pituitary hormone that stimulates not only somatic growth rate (Reinecke *et al.,* 2005; Canosa *et al.,* 2007) but also maturation, and also mediates osmoregulation and the stress response (Le Gac *et al.,* 1993; Wendelaar Bonga, 1997; Rousseau & Dufour, 2007).

We expected pituitary mRNA GH levels to be altered in parallel with body-size and maturation response to selection in the Large medaka line. However, pituitary mRNA GH levels were similar in the Large and Control lines. Instead, pituitary GH expression increased marginally significantly in the Small medaka line, which body size did not respond to selection. Specifically, the increase in GH was marginally significant in males only (+0.450, Model 8 in Appendix 4) but was of a similar amplitude in females (+0.448, results not shown). This counter-intuitive result may, in fact, be explained by the pleiotropic effects of GH on both somatic growth and maturation. In the Large medaka line, evolution towards faster somatic growth was possibly mediated by increased pituitary production of GH but, at the same time, evolution towards delayed maturation was possibly sustained by decreased pituitary GH production. The net result was that pituitary GH production was not significantly increased in the Large line compared to the Control line.

In contrast, in the Small medaka line the absence of body size evolution did not counteract evolution towards an increased pituitary production of GH, which was possibly associated with an increased reproductive investment. This hypothesis is supported by both results from the maturity probability model, in which the slope of the age effect on maturity probability was marginally significantly less negative in the Small compared to the Control line (Model 2, Appendix 4), and by increased egg size in the Small medaka line. Anyway, these effects in the Small line were weak, and further studies are needed to test whether reproductive traits do respond to selection for a smaller body size in the medaka.

Together with GH, we measured pituitary mRNA levels of the β subunits of the gonadotropins, the luteinizing (LHβ) and follicle-stimulating (FSHβ) hormones, which are known to stimulate steroidogenesis and gametogenesis and are involved in the onset of puberty in teleosts as in other vertebrates (Zohar *et al.,* 2010). We could not detect any significant effect of selection on pituitary gonadotropins in either the Large or Small medaka lines, suggesting than LHβ and FSHβ are less critical than GH to the evolution of life-history traits in the medaka. Interestingly, however, pituitary activity of the somatotropic (GH) and gonadotropic (LHβ and FSHβ) axes were highly positively correlated, suggesting that they are synergistic in their effects on medaka development. Similar results were previously found in the rainbow trout *Oncorhynchus mykiss* (Gomez *et al.,* 1999). Finally, the positive LH-GH correlation significantly increased in the Large medaka line, indicating that size-dependent selection may alter patterns of hormonal synergies. Future transcriptomic approaches on central and peripheral tissues will maybe provide a deeper understanding of the molecular regulation of response to size-dependent selection in the medaka.

### Conclusions

Inability of medaka to respond to selection for a smaller body size is a warning signal that calls for increasing research efforts to assess life-history evolvability in wild populations. A crucial line of work in achieving this goal will consist in accurately measuring the multivariate components of selection that act on correlated life-history traits such as body size and maturity (Le Rouzic *et al.*, Under Review; Lande & Arnold, 1983), both in the wild and in laboratory experiments. The other key element of this effort will rely on developing diagnosis tools to evaluate potential for (and signature of) adaptive response to size-dependent, anthropogenic selection (Therkildsen *et al.,* 2019). In the future, comprehensive approaches melting wide-spectrum candidate genes, transcriptomics and genome scans of experimentally- and wild-selected populations will probably be needed to finely decipher the molecular architectures that regulate the adaptive evolution of life histories and that ultimately support the maintenance of biodiversity and ecosystem productivity.

## Data Accessibility Statement

The data and the R scripts used for the analysis are fully available and provided as supplementary material.

## Acknowledgments

We are grateful to Prof. Kiyoshi Naruse (NIBB, Okazaki, Japan) for his support in obtaining and maintaining medaka from Kiyosu. We are also thankful to Prof. Finn-Arne Weltzien for providing several primer sequences for our candidate genes. We thank the people who helped us at the laboratory: Beatriz Decencière, Julien Hirschinger, Alice Lamoureux, Alexandre Macé and Yohann Chauvier.

## Author contributions

EE, ALR, GM and SD designed the study. CR, DC, AM, SA maintained the fish and performed selection, breeding, phenotyping and data collection. CR and GM performed mRNA measurements. EE and ALR made final analyses and finalized paper writing.

## Ethical statement

The protocols used in this study were designed to minimize discomfort, distress and pain of animals, and were approved by the Darwin Ethical committee (case file #Ce5/2010/041). The committee also confirmed that our methods were performed in accordance with the relevant guidelines and regulations on animal research.

## APPENDIX 1

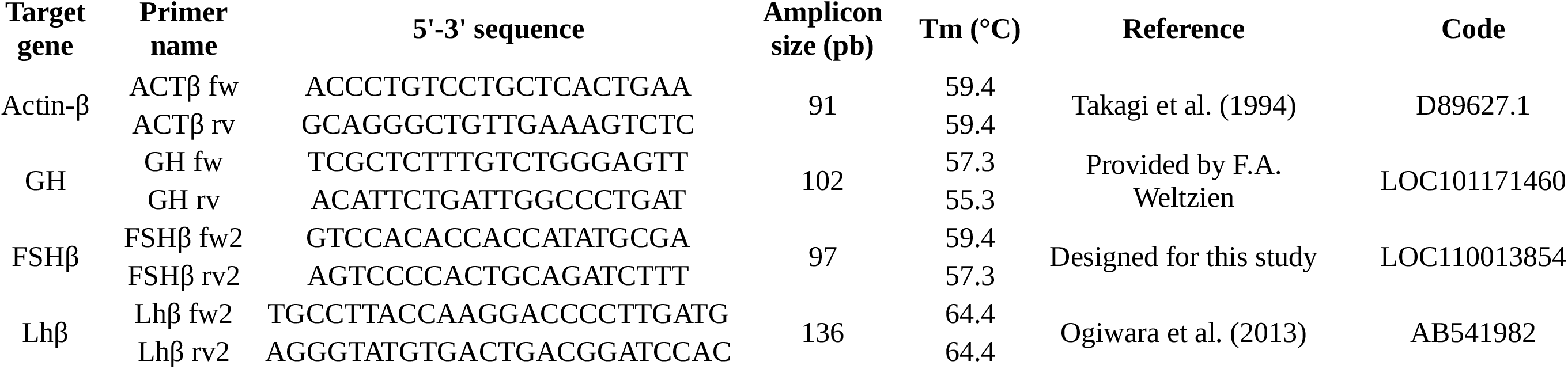
Primers for RTqPCR. Gene-specific primers (fw for forward and rv for reverse) were designed for amplification and quantification of mRNA of various medaka pituitary hormones by qRT-PCR using actin-β as a reference gene. GH: growth hormone, LHβ: luteinizing hormone β subunit, FSHβ follicle-stimulating hormone β subunit.

## APPENDIX 2

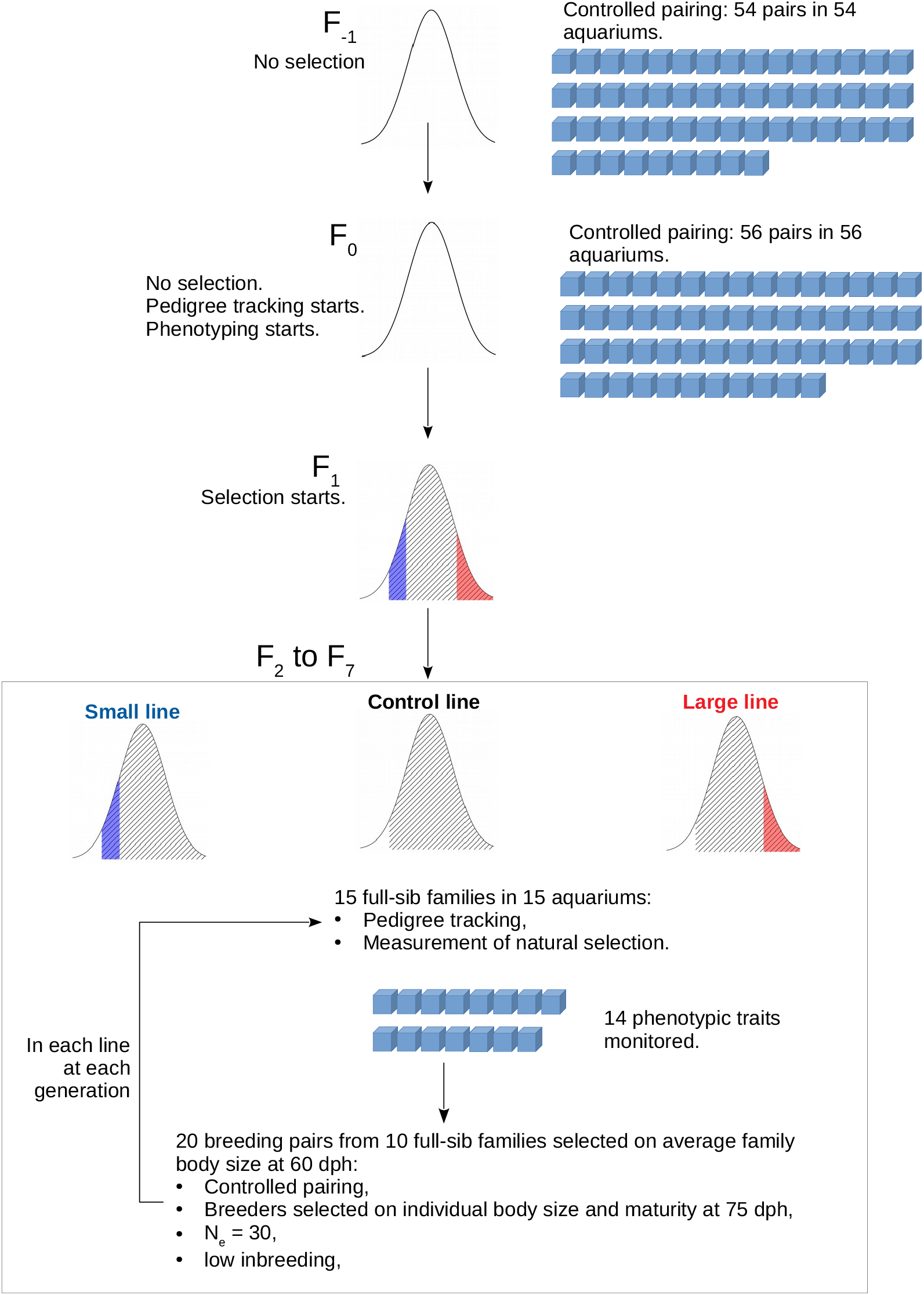
Schematic diagram of the experimental design. Curves show medaka standard body length distributions at 75 days-post-hatch (dph). Shaded areas represent mature individuals and coloured areas the mature individuals that were kept as breeders to form the next generation. Blue cubes represent the 3L aquariums in which medaka were maintained.

## APPENDIX 3

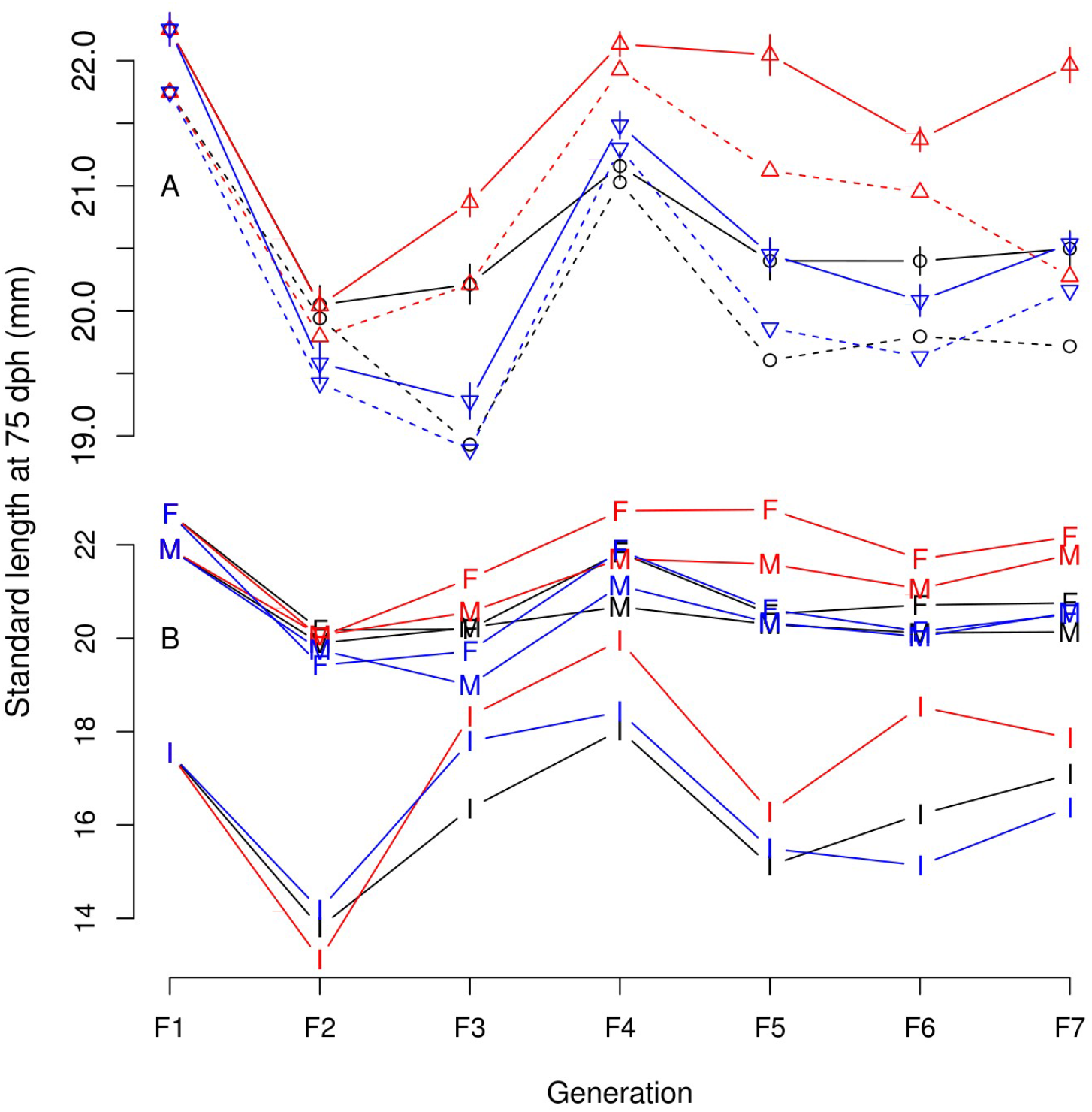
Raw medaka body-size time series response to bidirectional anthropogenic selection. A: mean standard body length of all (dashed lines) and mature fish (solid lines with ± SE) at 75 dph. Black circles: Control line, Blue bottom-pointing triangles: Small line, Red, top-pointing triangles: Large line. B: same as A but separately for immature (I), male (M) and female (F) fish and without error bars.

## APPENDIX 4

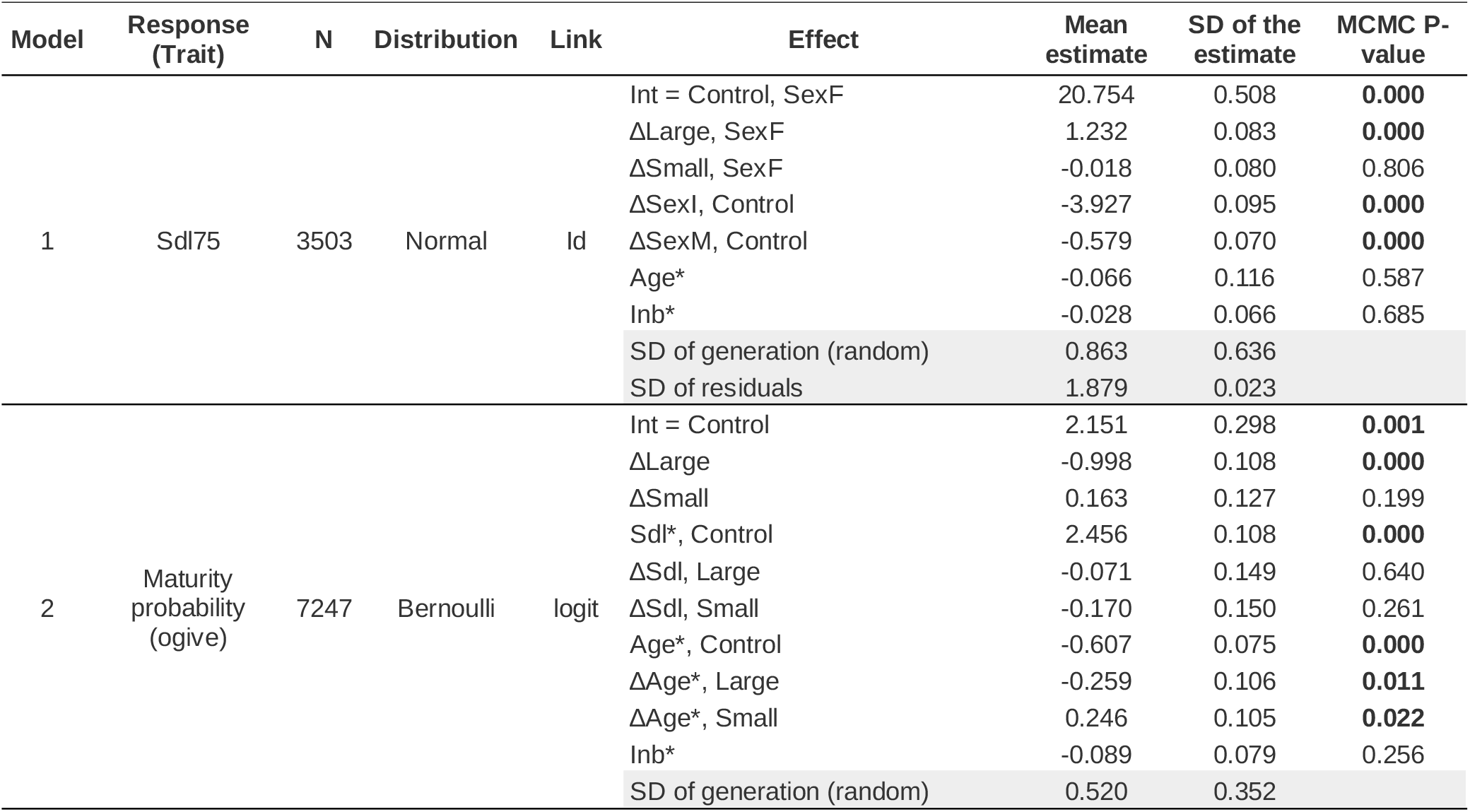
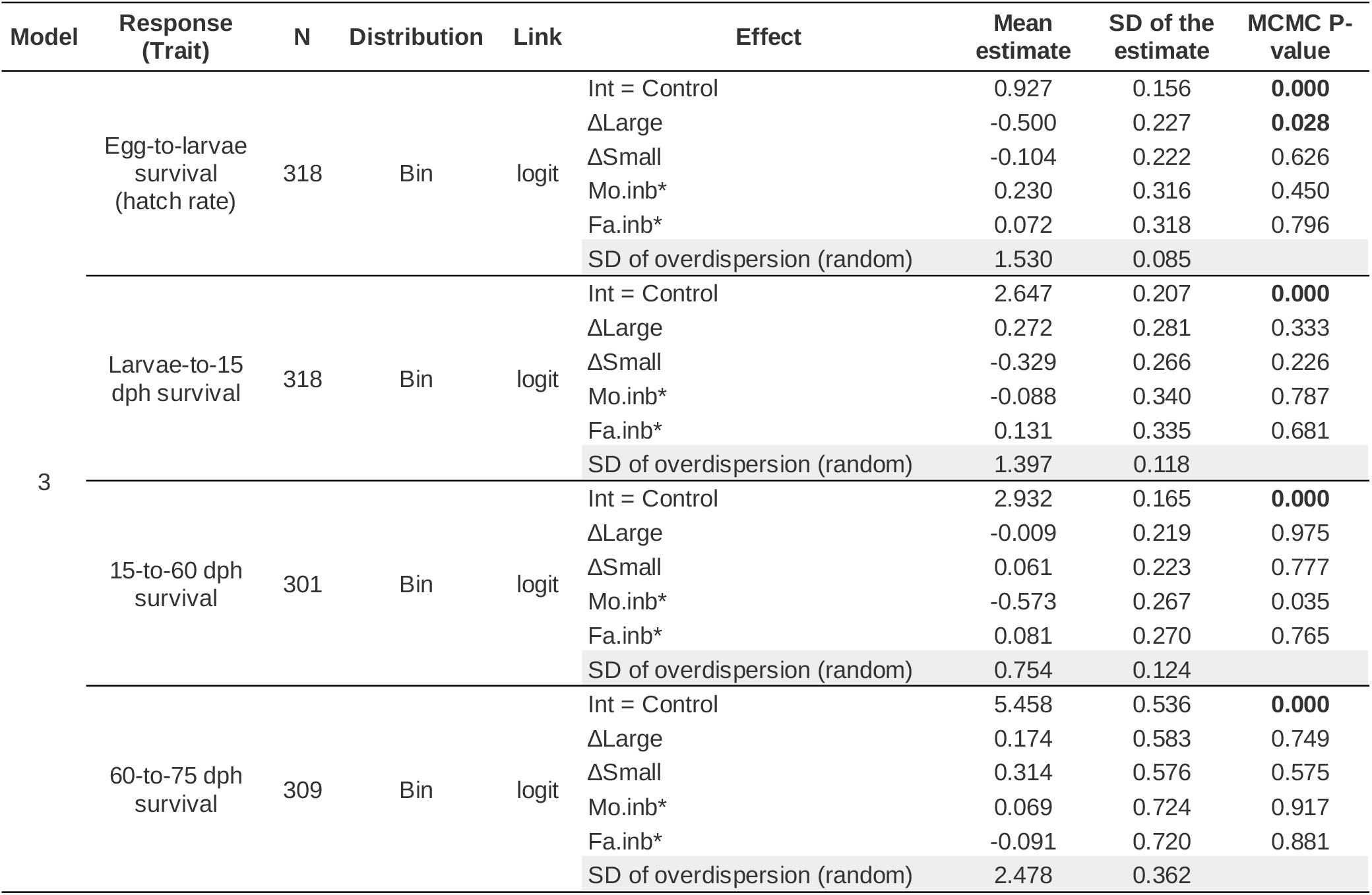
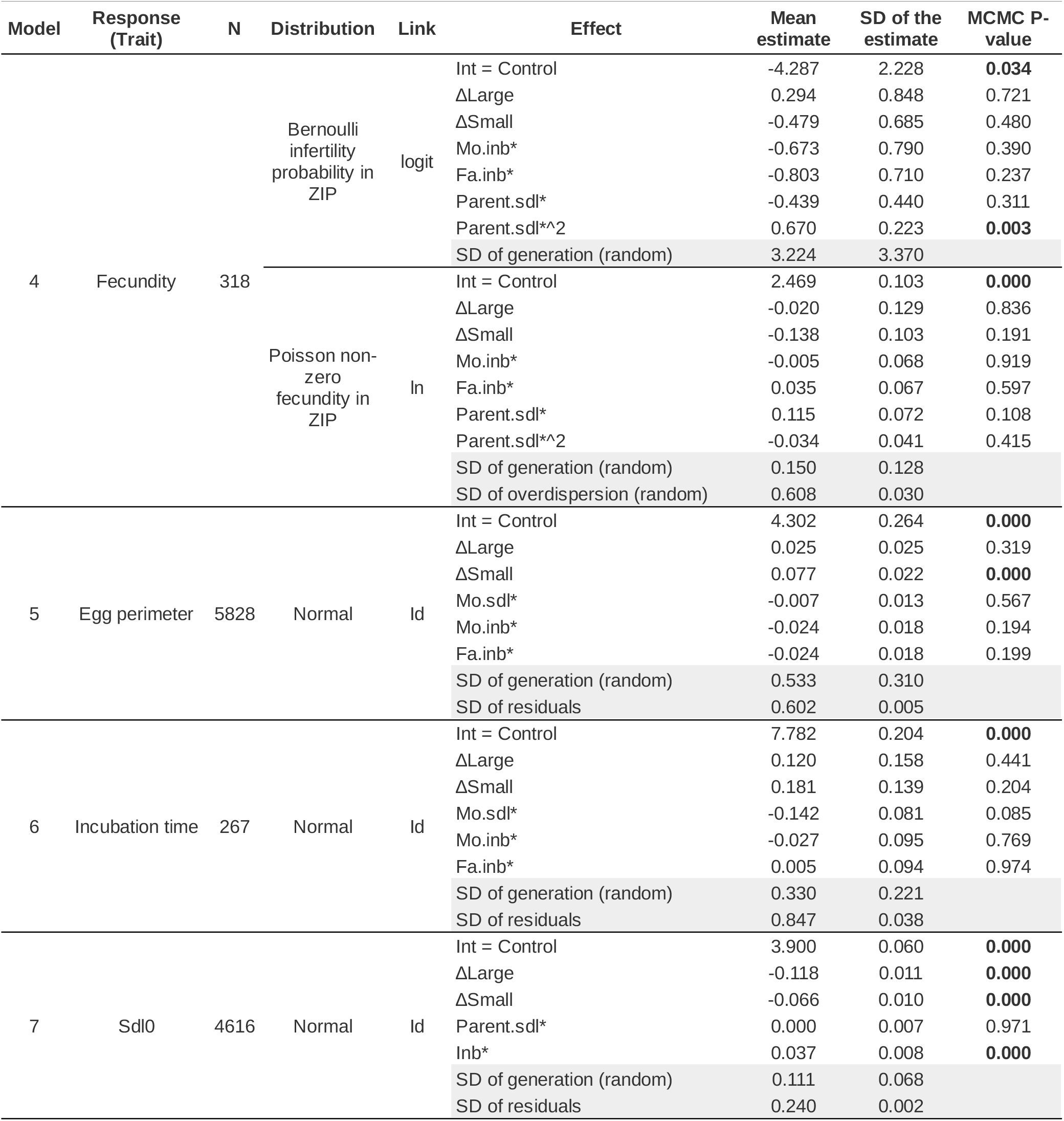
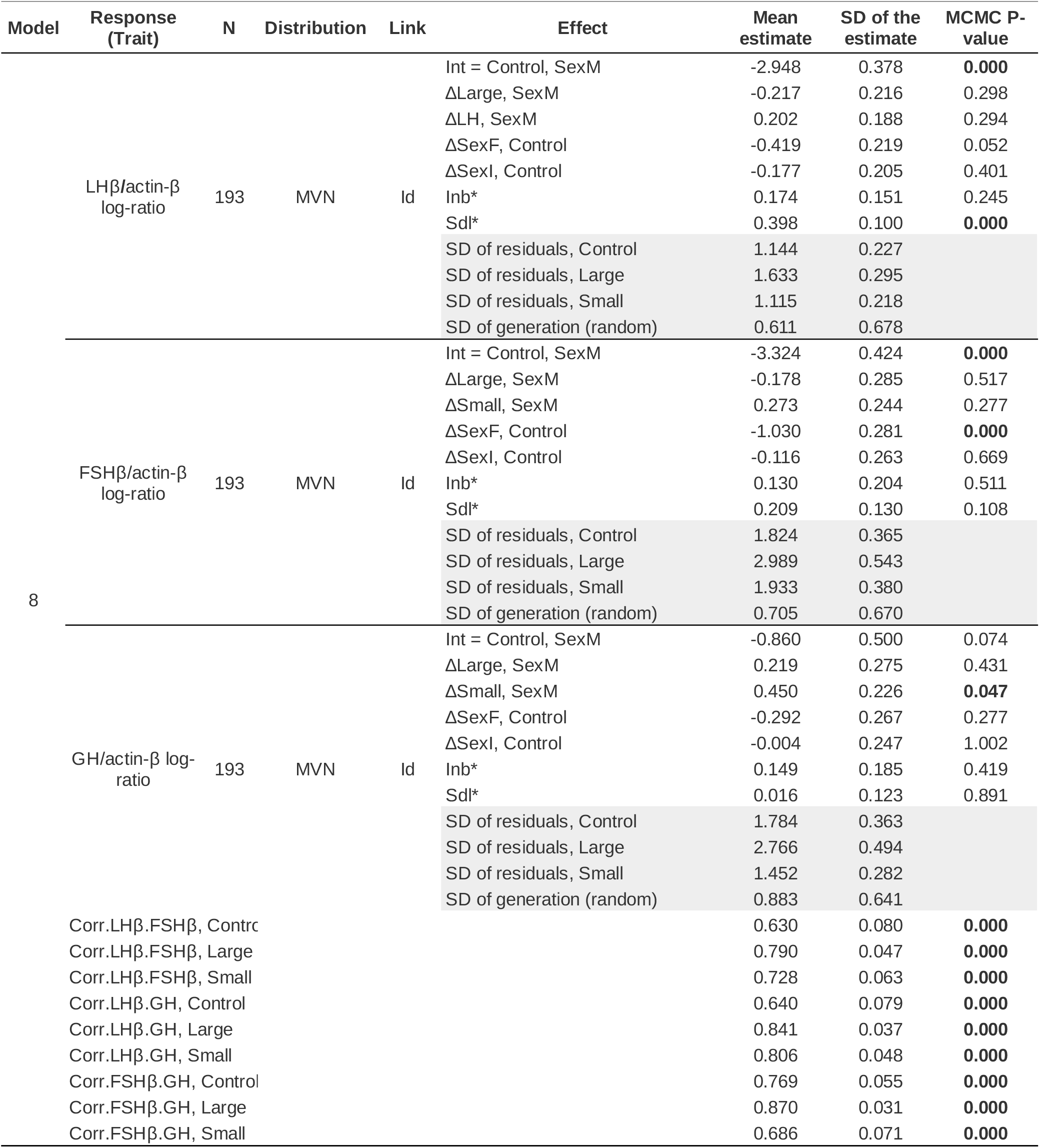
Summary of MCMC parameter estimates for models 1 to 8. We used an “effect” model parametrization in which the intercept is the mean value of the response variable (on the link scale) in the reference level of factor predictors. *ΔX* are estimates for the difference between mean value of the response variable in the factor level *X* and model intercept. Int: model intercept, F: female, I: immature, M: male, Control: random size-selected line, Large: line selected for a large body length, Small: line selected for a small body length, Inb: individual inbreeding coefficient, Sdl75: standard body length at 75 days post hatch, SD: standard deviation (shaded lines). * indicates that numeric predictors were scaled to zero mean and unity standard deviation (or 0.5 standard deviation in case of logistic regression following Gelman et al. 2008). ZIP in model 4 refers to the zero-inflated Poisson distribution. Corr.*X*.*Y* in model 8 is the correlation between pituitary expression levels of hormones *X* and *Y.*

| Model | Response (Trait) | N | Distribution | Link | Effect | Mean estimate | SD of the estimate | MCMC P-value |
| --- | --- | --- | --- | --- | --- | --- | --- | --- |
| 1 | Sdl75 | 3503 | Normal | Id | Int = Control, SexF | 20.754 | 0.508 | **0.000** |
| | | | | | ΔLarge, SexF | 1.232 | 0.083 | **0.000** |
| | | | | | ΔSmall, SexF | -0.018 | 0.080 | 0.806 |
| | | | | | ΔSexI, Control | -3.927 | 0.095 | **0.000** |
| | | | | | ΔSexM, Control | -0.579 | 0.070 | **0.000** |
|  |  |  |  |  | Age* | -0.066 | 0.116 | 0.587 |
|  |  |  |  |  | Inb* | -0.028 | 0.066 | 0.685 |
|  |  |  |  |  | SD of generation (random) | 0.863 | 0.636 |  |
|  |  |  |  |  | SD of residuals | 1.879 | 0.023 |  |
| 2 | Maturity probability (ogive) | 7247 | Bernoulli | logit | Int = Control | 2.151 | 0.298 | **0.001** |
| | | | | | ΔLarge | -0.998 | 0.108 | **0.000** |
| | | | | | ΔSmall | 0.163 | 0.127 | 0.199 |
|  |  |  |  |  | Sdl*, Control | 2.456 | 0.108 | **0.000** |
| | | | | | ΔSdl, Large | -0.071 | 0.149 | 0.640 |
| | | | | | ΔSdl, Small | -0.170 | 0.150 | 0.261 |
|  |  |  |  |  | Age*, Control | -0.607 | 0.075 | **0.000** |
| | | | | | ΔAge*, Large | -0.259 | 0.106 | **0.011** |
| | | | | | ΔAge*, Small | 0.246 | 0.105 | **0.022** |
|  |  |  |  |  | Inb* | -0.089 | 0.079 | 0.256 |
|  |  |  |  |  | SD of generation (random) | 0.520 | 0.352 |  |
| 3 | Egg-to-larvae survival (hatch rate) | 318 | Bin | logit | Int = Control | 0.927 | 0.156 | **0.000** |
| | | | | | ΔLarge | -0.500 | 0.227 | **0.028** |
| | | | | | ΔSmall | -0.104 | 0.222 | 0.626 |
|  |  |  |  |  | Mo.inb* | 0.230 | 0.316 | 0.450 |
|  |  |  |  |  | Fa.inb* | 0.072 | 0.318 | 0.796 |
|  |  |  |  |  | SD of overdispersion (random) | 1.530 | 0.085 |  |
|  | Larvae-to-15 dph survival | 318 | Bin | logit | Int = Control | 2.647 | 0.207 | **0.000** |
| | | | | | ΔLarge | 0.272 | 0.281 | 0.333 |
| | | | | | ΔSmall | -0.329 | 0.266 | 0.226 |
|  |  |  |  |  | Mo.inb* | -0.088 | 0.340 | 0.787 |
|  |  |  |  |  | Fa.inb* | 0.131 | 0.335 | 0.681 |
|  |  |  |  |  | SD of overdispersion (random) | 1.397 | 0.118 |  |
|  | 15-to-60 dph survival | 301 | Bin | logit | Int = Control | 2.932 | 0.165 | **0.000** |
| | | | | | ΔLarge | -0.009 | 0.219 | 0.975 |
| | | | | | ΔSmall | 0.061 | 0.223 | 0.777 |
|  |  |  |  |  | Mo.inb* | -0.573 | 0.267 | 0.035 |
|  |  |  |  |  | Fa.inb* | 0.081 | 0.270 | 0.765 |
|  |  |  |  |  | SD of overdispersion (random) | 0.754 | 0.124 |  |
|  | 60-to-75 dph survival | 309 | Bin | logit | Int = Control | 5.458 | 0.536 | **0.000** |
| | | | | | ΔLarge | 0.174 | 0.583 | 0.749 |
| | | | | | ΔSmall | 0.314 | 0.576 | 0.575 |
|  |  |  |  |  | Mo.inb* | 0.069 | 0.724 | 0.917 |
|  |  |  |  |  | Fa.inb* | -0.091 | 0.720 | 0.881 |
|  |  |  |  |  | SD of overdispersion (random) | 2.478 | 0.362 |  |
| 4 | Fecundity | 318 | Bernoulli infertility probability in ZIP | logit | Int = Control | -4.287 | 2.228 | **0.034** |
| | | | | | ΔLarge | 0.294 | 0.848 | 0.721 |
| | | | | | ΔSmall | -0.479 | 0.685 | 0.480 |
|  |  |  |  |  | Mo.inb* | -0.673 | 0.790 | 0.390 |
|  |  |  |  |  | Fa.inb* | -0.803 | 0.710 | 0.237 |
|  |  |  |  |  | Parent.sdl* | -0.439 | 0.440 | 0.311 |
|  |  |  |  |  | Parent.sdl*^2 | 0.670 | 0.223 | **0.003** |
|  |  |  |  |  | SD of generation (random) | 3.224 | 3.370 |  |
|  |  |  | Poisson non-zero fecundity in ZIP | ln | Int = Control | 2.469 | 0.103 | **0.000** |
| | | | | | ΔLarge | -0.020 | 0.129 | 0.836 |
| | | | | | ΔSmall | -0.138 | 0.103 | 0.191 |
|  |  |  |  |  | Mo.inb* | -0.005 | 0.068 | 0.919 |
|  |  |  |  |  | Fa.inb* | 0.035 | 0.067 | 0.597 |
|  |  |  |  |  | Parent.sdl* | 0.115 | 0.072 | 0.108 |
|  |  |  |  |  | Parent.sdl*^2 | -0.034 | 0.041 | 0.415 |
|  |  |  |  |  | SD of generation (random) | 0.150 | 0.128 |  |
|  |  |  |  |  | SD of overdispersion (random) | 0.608 | 0.030 |  |
| 5 | Egg perimeter | 5828 | Normal | ld | Int = Control | 4.302 | 0.264 | **0.000** |
| | | | | | ΔLarge | 0.025 | 0.025 | 0.319 |
| | | | | | ΔSmall | 0.077 | 0.022 | **0.000** |
|  |  |  |  |  | Mo.sdl* | -0.007 | 0.013 | 0.567 |
|  |  |  |  |  | Mo.inb* | -0.024 | 0.018 | 0.194 |
|  |  |  |  |  | Fa.inb* | -0.024 | 0.018 | 0.199 |
|  |  |  |  |  | SD of generation (random) | 0.533 | 0.310 |  |
|  |  |  |  |  | SD of residuals | 0.602 | 0.005 |  |
| 6 | Incubation time | 267 | Normal | ld | Int = Control | 7.782 | 0.204 | **0.000** |
| | | | | | ΔLarge | 0.120 | 0.158 | 0.441 |
| | | | | | ΔSmall | 0.181 | 0.139 | 0.204 |
|  |  |  |  |  | Mo.sdl* | -0.142 | 0.081 | 0.085 |
|  |  |  |  |  | Mo.inb* | -0.027 | 0.095 | 0.769 |
|  |  |  |  |  | Fa.inb* | 0.005 | 0.094 | 0.974 |
|  |  |  |  |  | SD of generation (random) | 0.330 | 0.221 |  |
|  |  |  |  |  | SD of residuals | 0.847 | 0.038 |  |
| 7 | Sdl0 | 4616 | Normal | ld | Int = Control | 3.900 | 0.060 | **0.000** |
| | | | | | ΔLarge | -0.118 | 0.011 | **0.000** |
| | | | | | ΔSmall | -0.066 | 0.010 | **0.000** |
|  |  |  |  |  | Parent.sdl* | 0.000 | 0.007 | 0.971 |
|  |  |  |  |  | Inb* | 0.037 | 0.008 | **0.000** |
|  |  |  |  |  | SD of generation (random) | 0.111 | 0.068 |  |
|  |  |  |  |  | SD of residuals | 0.240 | 0.002 |  |
| 8 | LHβ/actin-β log-ratio | 193 | MVN | Id | Int = Control, SexM | -2.948 | 0.378 | **0.000** |
| | | | | | ΔLarge, SexM | -0.217 | 0.216 | 0.298 |
| | | | | | ΔLH, SexM | 0.202 | 0.188 | 0.294 |
| | | | | | ΔSexF, Control | -0.419 | 0.219 | 0.052 |
| | | | | | ΔSexI, Control | -0.177 | 0.205 | 0.401 |
|  |  |  |  |  | Inb* | 0.174 | 0.151 | 0.245 |
|  |  |  |  |  | Sdl* | 0.398 | 0.100 | **0.000** |
|  |  |  |  |  | SD of residuals, Control | 1.144 | 0.227 |  |
|  |  |  |  |  | SD of residuals, Large | 1.633 | 0.295 |  |
|  |  |  |  |  | SD of residuals, Small | 1.115 | 0.218 |  |
|  |  |  |  |  | SD of generation (random) | 0.611 | 0.678 |  |
| | FSHβ/actin-β log-ratio | 193 | MVN | Id | Int = Control, SexM | -3.324 | 0.424 | **0.000** |
| | | | | | ΔLarge, SexM | -0.178 | 0.285 | 0.517 |
| | | | | | ΔSmall, SexM | 0.273 | 0.244 | 0.277 |
| | | | | | ΔSexF, Control | -1.030 | 0.281 | **0.000** |
| | | | | | ΔSexI, Control | -0.116 | 0.263 | 0.669 |
|  |  |  |  |  | Inb* | 0.130 | 0.204 | 0.511 |
|  |  |  |  |  | Sdl* | 0.209 | 0.130 | 0.108 |
|  |  |  |  |  | SD of residuals, Control | 1.824 | 0.365 |  |
|  |  |  |  |  | SD of residuals, Large | 2.989 | 0.543 |  |
|  |  |  |  |  | SD of residuals, Small | 1.933 | 0.380 |  |
|  |  |  |  |  | SD of generation (random) | 0.705 | 0.670 |  |
| | GH/actin-β log-ratio | 193 | MVN | Id | Int = Control, SexM | -0.860 | 0.500 | 0.074 |
| | | | | | ΔLarge, SexM | 0.219 | 0.275 | 0.431 |
| | | | | | ΔSmall, SexM | 0.450 | 0.226 | **0.047** |
| | | | | | ΔSexF, Control | -0.292 | 0.267 | 0.277 |
| | | | | | ΔSexI, Control | -0.004 | 0.247 | 1.002 |
|  |  |  |  |  | Inb* | 0.149 | 0.185 | 0.419 |
|  |  |  |  |  | Sdl* | 0.016 | 0.123 | 0.891 |
|  |  |  |  |  | SD of residuals, Control | 1.784 | 0.363 |  |
|  |  |  |  |  | SD of residuals, Large | 2.766 | 0.494 |  |
|  |  |  |  |  | SD of residuals, Small | 1.452 | 0.282 |  |
|  |  |  |  |  | SD of generation (random) | 0.883 | 0.641 |  |
| | Corr.LHβ.FSHβ, Contrc | | | | | 0.630 | 0.080 | **0.000** |
| | Corr.LHβ.FSHβ, Large | | | | | 0.790 | 0.047 | **0.000** |
| | Corr.LHβ.FSHβ, Small | | | | | 0.728 | 0.063 | **0.000** |
| | Corr.LHβ.GH, Control | | | | | 0.640 | 0.079 | **0.000** |
| | Corr.LHβ.GH, Large | | | | | 0.841 | 0.037 | **0.000** |
| | Corr.LHβ.GH, Small | | | | | 0.806 | 0.048 | **0.000** |
| | Corr.FSHβ.GH, Control | | | | | 0.769 | 0.055 | **0.000** |
| | Corr.FSHβ.GH, Large | | | | | 0.870 | 0.031 | **0.000** |
| | Corr.FSHβ.GH, Small | | | | | 0.686 | 0.071 | **0.000** |

## APPENDIX 5

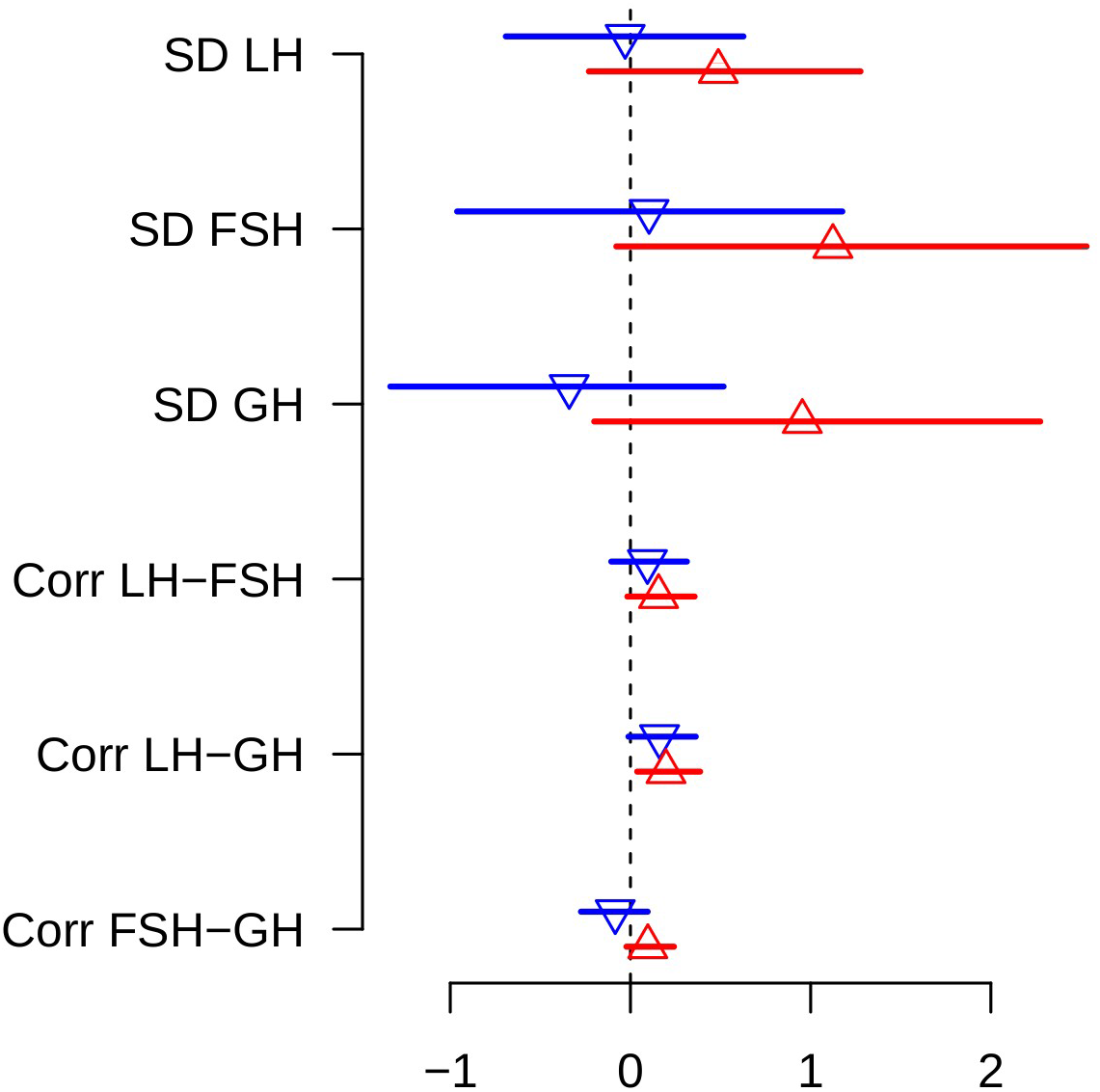
Residual standard deviations and correlations for pituitary hormone expression levels in size-selected medaka lines relative to the control line. Blue bottom-pointing triangles: Small line; Red, top-pointing triangles: Large line. SD: standard deviation, Corr: correlation, LH: pituitary gene expression of the β-subunit of the luteinizing hormone, FSH: pituitary gene expression of the β-subunit of the follicle-stimulating, GH: pituitary gene expression of the growth hormone.ç

## References

Amaral, I.P.G. & Johnston, I.A. 2012. Experimental selection for body size at age modifies early lifehistory traits and muscle gene expression in adult zebrafish. J. Exp. Biol. 215: 3895.

Barot, S., Heino, M., O’Brien, L. & Dieckmann, U. 2004. Estimating reaction norms for age and size at maturation when age at first reproduction is unknown. Evol. Ecol. Res. 6: 659–678.

Borrell, B. 2013. Ocean conservation: a big fight over little fish. Nature 493: 597–598.

Cameron, T.C., O’Sullivan, D., Reynolds, A., Piertney, S.B. & Benton, T.G. 2013. Eco-evolutionary dynamics in response to selection on life-history. Ecol. Lett. 16: 754–763.

Canosa, L.F., Chang, J.P. & Peter, R.E. 2007. Neuroendocrine control of growth hormone in fish. Gen. Comp. Endocrinol. 151: 1–26.

Carlson, S.M., Edeline, E., Vøllestad, L.A., Haugen, Thrond.O., Winfield, I.J., Fletcher, J.M., et al. 2007. Four decades of opposing natural and human-induced artificial selection acting on Windermere pike (Esox lucius). Ecol. Lett. 10: 512–521.

Conover, D.O. & Munch, S.B. 2002. Sustaining fisheries yields over evolutionary time scales. Science 297: 94–96.

Crow, J.F. & Kimura, M. 1970. An introduction to population genetics theory, 1st ed. Harper & Row, New York.

Devine, J.A. & Heino, M. 2011. Investigating the drivers of maturation dynamics in Barents Sea haddock (Melanogrammus aeglefinus). Fish. Res. 110: 441–449.

Diaz Pauli, B. & Heino, M. 2014. What can selection experiments teach us about fisheries-induced evolution? Biol. J. Linn. Soc. 111: 485–503.

Diaz Pauli, B., Kolding, J., Jeyakanth, G. & Heino, M. 2017. Effects of ambient oxygen and size-selective mortality on growth and maturation in guppies. Conserv. Physiol. 5: cox010–cox010.

Dieckmann, U. & Heino, M. 2007. Probabilistic maturation reaction norms: their history, strengths, and limitations. Mar. Ecol. Prog. Ser. 335: 253–269.

Duffy, T.A., Picha, M.E., Borski, R.J. & Conover, D.O. 2013. Circulating levels of plasma IGF-I during recovery from size-selective harvesting in Menidia menidia. Comp. Biochem. Physiol. A. Mol. Integr. Physiol. 166: 222–227.

Dunlop, E.S., Heino, M. & Dieckmann, U. 2009. Eco-genetic modeling of contemporary life-history evolution. Ecol. Appl. 19: 1815–1834.

Dunnington, E.A., Honaker, C.F., McGilliard, M.L. & Siegel, P.B. 2013. Phenotypic responses of chickens to long-term, bidirectional selection for juvenile body weight—Historical perspective. Poult. Sci. 92: 1724–1734.

Edeline, E. 2016. Life history evolution, human impacts on. In: The encyclopedia of evolutionary biology (R. Kliman, ed), pp. 335–342. Academic Press, Oxford.

Edeline, E., Carlson, S.M., Stige, L.C., Winfield, I.J., Fletcher, J.M., James, J.B., et al. 2007. Trait changes in a harvested population are driven by a dynamic tug-of-war between natural and harvest selection. Proc. Natl. Acad. Sci. 104: 15799–15804.

Edeline, E., Le Rouzic, A., Winfield, I.J., Fletcher, J.M., James, J.B., Stenseth, N.Chr., et al. 2009. Harvest-induced disruptive selection increases variance in fitness-related traits. Proc. R. Soc. Lond. B Biol. Sci. 276: 4163–4171.

Edley, M.T. & Law, R. 1988. Evolution of life histories and yields in experimental populations of Daphnia magna. Biol. J. Linn. Soc. 34: 309–326.

Eikeset, A.M., Dunlop, E.S., Heino, M., Storvik, G., Stenseth, N.C. & Dieckmann, U. 2016. Roles of density-dependent growth and life history evolution in accounting for fisheries-induced trait changes. Proc. Natl. Acad. Sci. 113: 15030–15035.

Falconer, D.S. 1973. Replicated selection for body weight in mice. Genet. Res. 22: 291–321.

Falconer, D.S. & Mackay, T.F.C. 1996. Introduction to quantitative genetics, 4th ed. Longman, Harlow, Essex, UK.

Fenberg, P.B. & Roy, K. 2008. Ecological and evolutionary consequences of size-selective harvesting: how much do we know? Mol. Ecol. 17: 209–220.

Gelman, A., Jakulin, A., Pittau, M.G. & Su, Y.-S. 2008. A weakly informative default prior distribution for logistic and other regression models. Ann. Appl. Stat. 2: 1360–1383.

Gelman, A., Meng, X.L. & Stern, H. 1996. Posterior predictive assessment of model fitness via realized discrepancies. Stat. Sin. 6: 733–807.

Gelman, A. & Rubin, D.B. 1992. Inference from iterative simulation using multiple sequences. Stat. Sci 457–472.

Gomez, J.M., Weil, C., Ollitrault, M., Le Bail, P.Y., Breton, B. & Le Gac, F. 1999. Growth hormone (GH) and gonadotropin subunit gene expression and pituitary and plasma changes during spermatogenesis and oogenesis in rainbow trout (Oncorhynchus mykiss). Gen. Comp. Endocrinol. 113: 413–428.

Hairston, N.G., Ellner, S.P., Geber, M.A., Yoshida, T. & Fox, J.A. 2005. Rapid evolution and the convergence of ecological and evolutionary time. Ecol. Lett. 8: 1114–1127.

Harney, E., Van Dooren, T.J.M., Paterson, S. & Plaistow, S.J. 2013. How to measure maturation: a comparison of probabilistic methods used to test for genotypic variation and plasticity in the decision to mature. Evolution 67: 525–538.

Heino, M., Díaz Pauli, B. & Dieckmann, U. 2015. Fisheries-induced evolution. Annu. Rev. Ecol. Evol. Syst. 46: 461–480.

Heino, M. & Dieckmann, U. 2008. Detecting fisheries-induced life-history evolution: an overview of the reaction-norm approach. Bull. Mar. Sci. 83: 69–93.

Heino, M., Dieckmann, U. & Godø, O.R. 2002. Measuring probabilistic reaction norms for age and size at maturation. Evolution 56: 669–678.

Hill, W.G. & Robertson, A. 1966. The effect of linkage on limits to artificial selection. Genet. Res. 8: 269–294.

Hillesheim, E. & Stearns, S.C. 1991. The responses of Drosophila melanogaster to artificial selection on body weight and its phenotypic plasticity in two larval food environments. Evolution 45: 1909–1923.

Kellner, K. 2019. jagsUI: a wrapper around “rjags” to streamline “JAGS” analyses.

Kendall, N.W., Hard, J.J. & Quinn, T.P. 2009. Quantifying six decades of fishery selection for size and age at maturity in sockeye salmon. Evol. Appl. 2: 523–536.

Kinoshita, M., Murata, K., Naruse, K. & Tanaka, M. 2009. Medaka. Biology, management and experimental protocols, 1st ed. Wiley, Ames (USA).

Koressaar, T. & Remm, M. 2007. Enhancements and modifications of primer design program Primer3. Bioinforma. Oxf. Engl. 23: 1289–1291.

Kuparinen, A., Kuikka, S. & Merilä, J. 2009. Estimating fisheries-induced selection: traditional gear selectivity research meets fisheries-induced evolution. Evol. Appl. 2: 234–243.

Kuparinen, A. & Merilä, J. 2007. Detecting and managing fisheries-induced evolution. Trends Ecol. Evol. 22: 652–659.

Lagler, K.F. 1968. Capture, sampling and examination of fishes. In: Methods for assessment of fish production in fresh waters (W. E. Ricker, ed), p. 313. Blackwell Publishing Ltd, Oxford.

Lande, R. & Arnold, S.J. 1983. The measurement of selection on correlated characters. Evolution 37: 1210–1226.

Law, R. 2000. Fishing, selection, and phenotypic evolution. ICES J. Mar. Sci. J. Cons. 57: 659–668.

Le Gac, F., Blaise, O., Fostier, A., Le Bail, P.Y., Loir, M., Mourot, B., et al. 1993. Growth hormone (GH) and reproduction: a review. Fish Physiol. Biochem. 11: 219–232.

Le Rouzic, A., Renneville, C., Millot, A., Agostini, S., Carmignac, D. & Edeline, E. Under Review. Unidirectional response to bidirectional selection on body size. II Quantitative genetics. Ecol. Evol.

Lunn, D., Jackson, C., Best, N., Thomas, A. & Spiegelhalter, D. 2012. The BUGS book: a practical introduction to Bayesian analysis, 1st ed. Chapman & Hall, Boca Raton.

Lynch, M. & Walsh, B. 2018. Evolution and selection of quantitative traits, 1st ed. Oxford University Press, New York.

Macarthur, J.W. 1949. Selection for small and large body size in the house mouse. Genetics 34: 194209.

Marty, L., Rochet, M.-J. & Ernande, B. 2014. Temporal trends in age and size at maturation of four North Sea gadid species: Cod, haddock, whiting and Norway pout. Mar. Ecol. Prog. Ser. 497.

Millar, R.B. & Fryer, R.J. 1999. Estimating the size-selection curves of towed gears, traps, nets and hooks. Rev. Fish Biol. Fish. 9: 89–116.

Morita, K. & Fukuwaka, M. 2006. Does size matter most? The effect of growth history on probabilistic reaction norm for salmon maturation. Evolution 60: 1516–1521.

Ogiwara, K., Fujimori, C., Rajapakse, S. & Takahashi, T. 2013. Characterization of luteinizing hormone and luteinizing hormone receptor and their indispensable role in the ovulatory process of the medaka. PLOS ONE 8: e54482.

Partridge, L., Langelan, R., Fowler, K., Zwaan, B. & French, V. 1999. Correlated responses to selection on body size in Drosophila melanogaster. Genet. Res. 74: 43–54.

Phillips, P.C., Whitlock, M.C. & Fowler, K. 2001. Inbreeding changes the shape of the genetic covariance matrix in Drosophila melanogaster. Genetics 158: 1137–1145.

Plummer, M. 2003. JAGS: a program for analysis of Bayesian graphical models using Gibbs sampling. Vienna, Austria.

R Core Team. 2020. R: a language and environment for statistical computing. R Foundation for Statistical Computing, Vienna, Austria.

Reinecke, M., Bjornsson, B.T., Dickhoff, W.W., McCormick, S.D., Navarro, I., Power, D.M., et al. 2005. Growth hormone and insulin-like growth factors in fish: where we are and where to go. Gen. Comp. Endocrinol. 142: 20–24.

Robertson, A. 1960. A theory of limits in artificial selection. Proc. R. Soc. Lond. B Biol. Sci. 153: 234–249.

Roff, D.A. 1992. The evolution of life histories, 1st ed. Chapman & Hall, New York.

Rousseau, K. & Dufour, S. 2007. Comparative aspects of GH and metabolic regulation in lower vertebrates. Neuroendocrinology 86: 165–174.

Silva, A., Faria, S. & Nunes, C. 2013. Long-term changes in maturation of sardine, Sardina pilchardus, in Portuguese waters. Sci. Mar. 77: 429–438.

Sinnwell, J.P., Therneau, T.M. & Schaid, D.J. 2014. The kinship2 R package for pedigree data. Hum. Hered. 78: 91–93.

Spivakov, M., Auer, T.O., Peravali, R., Dunham, I., Dolle, D., Fujiyama, A., et al. 2014. Genomic and phenotypic characterization of a wild medaka population: towards the establishment of an isogenic population genetic resource in fish. G3 Genes Genomes Genet. 4: 433–445.

Takagi, S., Sasado, T., Tamiya, G., Ozato, K., Wakamatsu, Y., Takeshita, A., et al. 1994. An efficient expression vector for transgenic medaka construction. Mol. Mar. Biol. Biotechnol. 3: 192–199.

Therkildsen, N.O., Wilder, A.P., Conover, D.O., Munch, S.B., Baumann, H. & Palumbi, S.R. 2019. Contrasting genomic shifts underlie parallel phenotypic evolution in response to fishing. Science 365: 487.

Trippel, E.A. 1995. Age at maturity as a stress indicator in fisheries. BioScience 45: 759–771.

Uusi-Heikkilä, S., Whiteley, A.R., Kuparinen, A., Matsumura, S., Venturelli, P.A., Wolter, C., et al. 2015. The evolutionary legacy of size-selective harvesting extends from genes to populations. Evol. Appl. 8: 597–620.

Van Dooren, T.J.M., Tully, T. & Ferrière, R. 2005. The analysis of reaction norms for age and size at maturity using maturation rate models. Evolution 59: 500–506.

van Wijk, S.J., Taylor, M.I., Creer, S., Dreyer, C., Rodrigues, F.M., Ramnarine, I.W., et al. 2013. Experimental harvesting of fish populations drives genetically based shifts in body size and maturation. Front. Ecol. Environ. 11: 181–187.

Walsh, M.R., Munch, S.B., Chiba, S. & Conover, D.O. 2006. Maladaptive changes in multiple traits caused by fishing: impediments to population recovery. Ecol. Lett. 9: 142–148.

Weber, K.E. & Diggins, L.T. 1990. Increased selection response in larger populations. II. Selection for ethanol vapor resistance in Drosophila melanogaster at two population sizes. Genetics 125: 585–597.

Wendelaar Bonga, S.E. 1997. The stress response in fish. Physiol. Rev. 77: 591–625.

Yamamoto, T. 1975. Medaka (killifish): biology and strains, 1st ed. Keigaku Pub. Co, Tokyo.

Zohar, Y., Munoz-Cueto, J.A., Elizur, A. & Kah, O. 2010. Neuroendocrinology of reproduction in teleost fish. Gen. Comp. Endocrinol. 165: 438–455.

