## Supplementary material for "Unidirectional response to bidirectional selection on body size. I. Phenotypic, life history and endocrine response": AgeCorrect.R

```
AgeCorrect <- function(sdl, Phenotyping="P3", Age=75, method=c("previous", "P0", "avg"), returnAge=FALSE) {
    # Correction methods:
    #    previous: corrects the sdl by calculating the growth between the phenotyping and the previous one.
    #    F0: computes the average growth from F0
    #    avg: computes the average effect of age around the phenotyping point throughout all the dataset
    
    stopifnot (Phenotyping != "P0")
    Phenotyping.previous <- "P0"
    if (method[1] == "previous") 
        Phenotyping.previous <- paste0("P", as.numeric(strsplit(Phenotyping, split="")[[1]][2])-1)
        
    sdl$Date <- as.Date(sdl$Date, format="%d/%m/%Y")
    
    # Timing events by family
    birth <- tapply(sdl$Date[sdl$Phenotyping=="P0"], INDEX=sdl$Family[sdl$Phenotyping=="P0"], FUN=mean)
    dateP <- tapply(sdl$Date[sdl$Phenotyping==Phenotyping], INDEX=sdl$Family[sdl$Phenotyping==Phenotyping], FUN=mean)
    dateP1 <- tapply(sdl$Date[sdl$Phenotyping==Phenotyping.previous], INDEX=sdl$Family[sdl$Phenotyping==Phenotyping.previous], FUN=mean)
    
    sdlP <- tapply(sdl$Sdl[sdl$Phenotyping==Phenotyping], INDEX=sdl$Family[sdl$Phenotyping==Phenotyping], FUN=mean)
    sdlP1 <- tapply(sdl$Sdl[sdl$Phenotyping==Phenotyping.previous], INDEX=sdl$Family[sdl$Phenotyping==Phenotyping.previous], FUN=mean)
    
    # for the avg method (regression by family -- should be virtually identical to the full regression
    avg.slope <- coef(lm(sdlP ~ I(dateP-birth)))[2]
    
    mapply(function(phenot, fam, sl) {
        fam <- as.character(fam)
        if (phenot != Phenotyping) {
            NA
        } else {
            if (returnAge) dateP[fam]-birth[fam]
            else {
                a <- if (method[1] == "avg") avg.slope
                    else (sdlP[fam]-sdlP1[fam])/(dateP[fam]-dateP1[fam])
                sl+a*(Age-(dateP[fam]-birth[fam]))
            }
        }}, sdl$Phenotyping, sdl$Family, sdl$Sdl)
}

Age <- function(sdl, FUN=median) {
    # Returns the age at which every phenotypic measurement was done
    
    sdl$Date <- as.Date(sdl$Date, format="%d/%m/%Y")
    
    # Timing events by family
    birth <- tapply(sdl$Date[sdl$Phenotyping=="P0"], INDEX=sdl$Family[sdl$Phenotyping=="P0"], FUN=FUN)
    dateP <- tapply(sdl$Date, INDEX=list(sdl$Family, sdl$Phenotyping), FUN=FUN)
    mapply(function(phenot, fam) {
        dateP[fam, phenot]-birth[fam]
    }, sdl$Phenotyping, sdl$Family)
}
```
