## Supplementary material for "Unidirectional response to bidirectional selection on body size. I. Phenotypic, life history and endocrine response": New2Old.R

```
################################# New2Old.R ##########################
#
# Manipulates new and old aquarium/family numbers
#
# Author: Arnaud Le Rouzic <>
# Version 1.3      2017-05-31
#
# Distributed under the WTFPL v2, no warranty
#
######################################################################


##### Summary
######### check ID validity
#
# is.ValidAqua(x)
# is.ValidID(x)
#
######### new vs old
#
# is.NewAqua(x)
# is.OldAqua(x)
# is.NewID(x)
# is.OldID(x)
#
######### change old to new and vice versa
#
# Aqua2New(x)
# Aqua2Old(x)
# ID2New(x)
# ID2Old(x)
#
########## decomposition
#
# decompAqua(x)
# decompID(x)
# 
# which provide lists with various elements, such as:
# ..$Generation
# ..$Line
# ..$Family
# etc (check the code for details)


############################ Aquarium IDs ####################

makenewID.aquaID <- function(idobj) {
	idobj$AquaNew <- Aqua2New(idobj$AquaOld)
	paste0(idobj$Generation, ".", idobj$AquaNew)
}

makeoldID.aquaID <- function(idobj) {
	idobj$AquaOld <- Aqua2Old(idobj$AquaNew)
	paste0(idobj$Generation, idobj$AquaOld)
}
 
decompOldAqua <- function(old.aqua) {
    stopifnot(is.OldAqua(old.aqua))
	rr <- regexec(pattern="^(F\\d)[.](\\d+)([A-Z])$", old.aqua)
	res <- regmatches(old.aqua, rr)[[1]]
    
    generation <- res[2]
    oldfam     <- as.numeric(res[3])
    oldrep     <- res[4]

    line       <- "P"
    if (oldfam > 20 && oldfam <= 40) line <- "G"
    if (oldfam > 40) line <- "A"
    newfam <- oldfam
    if (line=="G") newfam <- newfam - 20
    if (line=="A") newfam <- newfam - 40
	
    ans <- list(
        Generation = generation, 
        Family     = as.character(oldfam), 
        FamilyFull = paste0(generation, ".", as.character(oldfam)),
        FamilyNew  = paste0(newfam, line),
        FamilyOld  = as.character(oldfam), 
        Aqua       = paste0(oldfam, oldrep),
        AquaFull   = paste0(generation, ".", oldfam, oldrep),
        AquaNew    = paste0(newfam, line, tolower(oldrep)), 
        AquaOld    = paste0(oldfam, oldrep), 
        Line       = line)
    class(ans) <- c("aquaID", class(ans))
    ans
}

decompNewAqua <- function(new.aqua) {
    stopifnot(is.NewAqua(new.aqua))
	rr <- regexec(pattern="^(F\\d+)[.](\\d+)([PGA0])([a-z])$", new.aqua)
	res <- regmatches(new.aqua, rr)[[1]]
    
    generation <- res[2]
    newfam     <- paste0(res[3], res[4])
    newrep     <- res[5]
    line       <- res[4]
    
    oldfam <- as.numeric(res[3])
    if (line == "G") oldfam <- oldfam + 20
    if (line == "A") oldfam <- oldfam + 40   
     
	ans <- list(
        Generation = generation, 
        Family     = newfam, 
        FamilyFull = paste0(generation, ".", newfam),
        FamilyNew  = newfam,
        FamilyOld  = as.character(oldfam), 
        Aqua       = paste0(newfam, newrep),
        AquaFull   = paste0(generation, ".", newfam, newrep),
        AquaNew    = paste0(newfam, newrep), 
        AquaOld    = paste0(oldfam, toupper(newrep)),  
        Line       = line)
    class(ans) <- c("aquaID", class(ans))
    ans
}
    
is.ValidAqua <- function(aqua) {
    is.NewAqua(aqua) | is.OldAqua(aqua)
}

is.NewAqua <- function(aqua) {
    grepl(pattern="^F\\d+.\\d+[OAGP][a-z]$", aqua)
}  

is.OldAqua <- function(aqua) {
    grepl(pattern="^F\\d+.\\d+[A-Z]$", aqua)   
}  

decompAqua <- function(aqua) {
    ans <- lapply(aqua, function(aqua.u) {
        if (is.NewAqua(aqua.u)) decompNewAqua(aqua.u)
        else if (is.OldAqua(aqua.u)) decompOldAqua(aqua.u)
        else NA })
    if (length(ans) == 1) {ans[[1]]} else {ans}
}

old2newAqua.u <- function(old.u) {
    stopifnot(is.ValidAqua(old.u))
    if (is.NewAqua(old.u)) return(old.u)
	aquaobj <- decompOldAqua(old.u)
	makenewAqua(idobj)
}

new2oldAqua.u <- function(new.u) {
    stopifnot(is.ValidAqua(new.u))
    if (is.OldAqua(new.u)) return(new.u)
	aquaobj <- decompNewAqua(new.u)
	makeoldAqua(aquaobj)
}

Aqua2New <- function(oldaq) {
	sapply(oldaq, Aqua2New.u)
}

Aqua2Old <- function(newaq) {
	sapply(newaq, Aqua2Old.u)
}

#################################### Fish IDs ##########################


makenewID.fishID <- function(idobj) {
	idobj$AquaNew <- Aqua2New(idobj$AquaOld)
	paste0(idobj$Generation, ".", idobj$AquaNew, ".", idobj$Sex, idobj$BreederIndex)
}

makeoldID.fishID <- function(idobj) {
	idobj$AquaOld <- Aqua2Old(idobj$AquaNew)
	paste0(idobj$Generation, idobj$AquaOld, idobj$Sex, idobj$BreederIndex, idobj$Line)
}

decompOldID <- function(old.id) {
    stopifnot(is.OldID(old.id))
	rr <- regexec(pattern="^(F\\d)(\\d+)([A-Z])([FMI])(\\d+)([OAGP])$", old.id)
	res <- regmatches(old.id, rr)[[1]]
    
    generation <- res[2]
    oldfam     <- as.numeric(res[3])
    oldrep     <- res[4]
    sex        <- res[5]
    index      <- res[6]
    line       <- res[7]
    newfam <- oldfam
    if (line=="G") newfam <- newfam - 20
    if (line=="A") newfam <- newfam - 40
	ans <- list(
        Generation = generation, 
        Family     = as.character(oldfam), 
        FamilyFull = paste0(generation, ".", as.character(oldfam)),    
        FamilyNew  = paste0(newfam, line),
        FamilyOld  = as.character(oldfam), 
        Aqua       = paste0(oldfam, oldrep),
        AquaFull   = paste0(generation, ".", oldfam, oldrep),                
        AquaNew    = paste0(newfam, line, tolower(oldrep)), 
        AquaOld    = paste0(oldfam, oldrep), 
        Sex        = sex, 
        BreederIndex=index, 
        Line       = line)
    class(ans) <- c("fishID", class(ans))
    ans
}

decompNewID <- function(new.id) {
    stopifnot(is.NewID(new.id))
	rr <- regexec(pattern="^(F\\d+)\\.(\\d+)([OAGP])([a-z])\\.([FMI])(\\d+)$", new.id)
	res <- regmatches(new.id, rr)[[1]]

    generation <- res[2]
    newfam     <- paste0(res[3], res[4])
    newrep     <- res[5]
    sex        <- res[6]
    index      <- res[7]
    line       <- res[4]
    
    oldfam <- as.numeric(res[3])
    if (line == "G") oldfam <- oldfam + 20
    if (line == "A") oldfam <- oldfam + 40   
	ans <- list(
        Generation = generation, 
        Family     = newfam, 
        FamilyFull = paste0(generation, ".", newfam),
        FamilyNew  = newfam,
        FamilyOld  = as.character(oldfam), 
        Aqua       = paste0(newfam, newrep),
        AquaFull   = paste0(generation, ".", newfam, newrep),        
        AquaNew    = paste0(newfam, newrep), 
        AquaOld    = paste0(oldfam, toupper(newrep)), 
        Sex        = sex, 
        BreederIndex=index, 
        Line       = line)
    class(ans) <- c("fishID", class(ans))
    ans
}

is.NewID <- function(id) {
    grepl(pattern="^F\\d+.\\d+[OAGP][abcd].[MFI]\\d+$", id)
}

is.OldID <- function(id) {
    grepl(pattern="^F\\d+[ABCD][MFI]\\d+[OAGP]$", id)
}

is.ValidID <- function(id) {
    is.NewID(id) | is.OldID(id)
}

decompID <- function(id) {
    ans <- lapply(id, function(id.u) {
        if (is.NewID(id.u)) decompNewID(id.u)
        else if (is.OldID(id.u)) decompOldID(id.u)
        else NA })
    if (length(ans) == 1) {ans[[1]]} else {ans}
}

old2newID.u <- function(old.u) {
    stopifnot(is.ValidID(old.u))
    if (is.NewID(old.u)) return(old.u)
	idobj <- decompOldID(old.u)
	makenewID(idobj)
}

new2oldID.u <- function(new.u) {
    stopifnot(is.ValidID(new.u))
    if (is.OldID(new.u)) return(new.u)
	idobj <- decompNewID(new.u)
	makeoldID(idobj)
}

ID2New <- function(oldid) {
	sapply(oldid, old2newID.u)
}

ID2Old <- function(newid) {
	sapply(newid, new2oldID.u)
}
```
