## Supplementary material for "Unidirectional response to bidirectional selection on body size. I. Phenotypic, life history and endocrine response": Eggs_size.R

```
#==========================================================================================================================================================
#			R script for Unidirectional response to bidirectional selection on body size. I. Phenotypic, life history and endocrine response.
#			Eric Edeline July 2020
#
#			JAGS MODEL FOR EGG PERIMETER -- EFFECT PARAMETRIZATION
#
#			/!\ BEFORE RUNNING:	SET WORKING DIRECTORY TO THE Data_and_code FOLDER ON YOUR OWN MACHINE
#						FOR INSTANCE ON MY MACHINE: setwd("~/Dropbox/Article_Clem/Manuscript/Submission/Ecology_and_Evolution/Data_and_codes")
#==========================================================================================================================================================
jags_model = "
  model {
### Likelihood
    for (i in 1:N) {
      y[i] ~ dnorm(mu[i], 1/(sigma^2))
      mu[i] <- alpha0[Gener[i]] + beta0 + beta1[Line[i]] + beta2  * Mo.Sdl[i] + beta3 * Mo.Inb[i] + beta4 * Fa.Inb[i]
    }
### Priors:
  # Residuals
  sigma ~ dunif(0, 10)
  # Random mother effect
  for(i in 1:ngener){
    alpha0[i] ~ dnorm(0.0, 1/(sigma.alpha0^2))  
  }
  sigma.alpha0 ~ dunif(0.0, 10)
  # Fixed effects
  beta0 ~ dnorm(0.0, 0.001)		# Intercept = line A at an average Mo.Sdl, Mo.Inb and Fa.Inb
  beta1[1] <- 0.0			# Constraint for the effect parametrization
  beta1[2] ~ dnorm(0.0 , 0.001)  	# Effect of G line on intercept
  beta1[3] ~ dnorm(0.0 , 0.001)		# Effect of P line on intercept
  beta2 ~ dnorm(0.0, 0.001)		# Slope of Mo.Sdl effect
  beta3 ~ dnorm(0.0, 0.001)		# Slope of Mo.Inb effect
  beta4 ~ dnorm(0.0, 0.001)		# Slope of Fa.Inb effect
# Assess model fit using sums-of-squares type of discrepancy
  for(i in 1:N){
    sq[i] <- pow(y[i] - mu[i], 2)			#squared residuals
    # Generate replicate data and compute fit stats for them
    y.new[i] ~ dnorm(mu[i], 1/sigma^2)			#ideal data produced from model parameters
    sq.new[i] <- pow(y.new[i] - mu[i], 2)	#squared residuals from ideal data
    }
    fit <- sum(sq[])			#sum of squared residuals from true data
    fit.new <- sum(sq.new[])		#sum of squared residuals from ideal data
    test <- step(fit.new - fit)		#test whether new data set more extreme
    bpvalue <- mean(test)		#Bayesian p-value
    c.hat <- fit/(fit.new + 0.00001)	# lack-of-fit ratio (moderate for c-hat up to 4, Mazerolles 2015 in AHM p. 252)
} "
writeLines( jags_model , con = "./Scripts/Egg_size/Normal_Eggs.jags" )

#===========================================================================================================================================================
#	PARAMETERS TO MONITOR
#===========================================================================================================================================================
parameters = c("sigma", "sigma.alpha0", "beta0", "beta1", "beta2", "beta3", "beta4", "bpvalue", "c.hat")

#===========================================================================================================================================================
#	DATA IMPORT AND PREPARATION
#===========================================================================================================================================================
### Import data on eggs
Data <- read.table(file = "./Data/Egg_sizes.txt", header = TRUE, sep = "\t", dec = ".", na.strings = "NA")
### Keep only egg perimeter metric
Data <- Data[,c(1,5,22:24)]
### Import data on families
Fam <- read.table(file = "./Data/Family.txt", header = TRUE, sep = "\t", dec = ".", na.strings = "NA")
### Merge
Data = merge(Data, Fam, by.x="Family", by.y="Family")
### Keep only F3-to-F7 data
Data <- Data[Data$Generation.x != "F1" & Data$Generation.x != "F2",];
Data$Generation.x <- factor(Data$Generation.x); Data$Line <- factor(Data$Line); Data$Mother <- factor(Data$Mother); 
### Remove NA predictors
Data <- Data[!is.na(Data$Mothersdl),]

### Bundle data
jags.data <- list(N = length(Data$Perim.), y = Data$Perim., Gener = as.integer(Data$Generation.x), ngener = length(levels(Data$Generation.x)), Mother = as.integer(Data$Mother), nmother= length(levels(Data$Mother)), Line = as.integer(Data$Line), Mo.Sdl = (Data$Mothersdl - mean(Data$Mothersdl)) / sd(Data$Mothersdl), Mo.Inb = (Data$Mother.inb - mean(Data$Mother.inb)) / sd(Data$Mother.inb), Fa.Inb = (Data$Father.inb - mean(Data$Father.inb)) / sd(Data$Father.inb))

#===========================================================================================================================================================
#	MCMC SETTINGS
#===========================================================================================================================================================
ni <- 6000
nt <- 1
nb <- 5000
nc <- 3

#######################################################################################################################################
#	SAMPLING POSTERIOR
#######################################################################################################################################
library(jagsUI)
mod <- jags(data = jags.data, parameters.to.save = parameters, model.file = "./Scripts/Egg_size/Normal_Eggs.jags", n.chains = nc, n.thin = nt, n.iter = ni, n.burnin = nb, n.adapt = 2000, parallel = TRUE) #
```
