## Supplementary material for "Unidirectional response to bidirectional selection on body size. I. Phenotypic, life history and endocrine response": Fecundity.R

```
#==========================================================================================================================================================
#			R script for Unidirectional response to bidirectional selection on body size. I. Phenotypic, life history and endocrine response.
#			Eric Edeline July 2020
#
#			DATA MODELLING USING A ZERO-INFLATED POISSON MODEL -- EFFECT PARAMETERIZATION
#
#			/!\ BEFORE RUNNING:	SET WORKING DIRECTORY TO THE Data_and_code FOLDER ON YOUR OWN MACHINE
#						FOR INSTANCE ON MY MACHINE: setwd("~/Dropbox/Article_Clem/Manuscript/Submission/Ecology_and_Evolution/Data_and_codes")
#==========================================================================================================================================================
jags_model = "
  model {
#### Likelihood
    for (i in 1:N) {
	C[i] ~ dpois(lambda.eff[i])
	lambda.eff[i] <- lambda[i] * (1 - u[i])						# Composite Poisson mean
    	log(lambda[i]) <- offset[i] + alpha0[Gener[i]] + alpha1 + alpha2[Line[i]] + alpha3 * Mother.inb[i] + alpha4 * Father.inb[i] + alpha5 * Parent.sdl[i] + alpha6 * pow(Parent.sdl[i], 2) + eps[i]	# Poisson model
	u[i] ~ dbern(psi[i])
	logit(psi[i]) <- beta0[Gener[i]] + beta1 + beta2[Line[i]] + beta3 * Mother.inb[i] + beta4 * Father.inb[i] + beta5 * Parent.sdl[i] + beta6 * pow(Parent.sdl[i], 2)	# Bernoulli model, same linear predictor
	eps[i] ~ dnorm(0, tau.eps)
    }
#### Priors:
# Overdispersion parameter
    tau.eps <- pow(sigma.eps, -2)
    sigma.eps ~ dunif(0, 100)
# Random generation effects
    for(i in 1:ngener){
	alpha0[i] ~ dnorm(0, tau.alpha)
	beta0[i] ~ dnorm(0, tau.beta)
    }
    tau.alpha <- 1 / (sigma.alpha^2)
    sigma.alpha ~ dunif(0, 100)
    tau.beta <- 1 / (sigma.beta^2)
    sigma.beta ~ dunif(0, 100)
# Fixed effects, log regression
    alpha1 ~ dnorm(0, 0.001)	# Model intercept = log(non-zero fecundity) in A line
    alpha2[1] <- 0
    alpha2[2] ~ dnorm(0, 0.001)	# Effect of G line compared to A line
    alpha2[3] ~ dnorm(0, 0.001)	# Effect of P line compared to A line
    alpha3 ~ dnorm(0, 0.001)	# Slope of mother inbreeding effect
    alpha4 ~ dnorm(0, 0.001)	# Slope of father inbreeding effect
    alpha5 ~ dnorm(0, 0.001)	# Slope of body length effect
    alpha6 ~ dnorm(0, 0.001)	# Slope of squared body length effect
# Fixed effects, logit regression
    beta1 ~ dnorm(0, 0.001)	# Model intercept = logit(probability to spawn) in A line
    beta2[1] <- 0
    beta2[2] ~ dnorm(0, 0.001)	# Effect of G line compared to A line
    beta2[3] ~ dnorm(0, 0.001)	# Effect of P line compared to A line
    beta3 ~ dnorm(0, 0.001)	# Slope of mother inbreeding effect
    beta4 ~ dnorm(0, 0.001)	# Slope of father inbreeding effect
    beta5 ~ dnorm(0, 0.001)	# Slope of body length effect
    beta6 ~ dnorm(0, 0.001)	# Slope of squared body length effect

#### Assess model fit using sums-of-squares type of discrepancy
  for(i in 1:N){
    res[i] <- (C[i] - lambda.eff[i]) / sqrt(lambda.eff[i] + 0.0001)			# Pearson residuals
    # Generate replicate data and compute fit stats for them
    C.new[i] ~ dpois(lambda.eff[i])						# Ideal data produced from model parameters
    res.new[i] <- (C.new[i] - lambda.eff[i]) / sqrt(lambda.eff[i] + 0.0001)		# Pearson residuals from ideal data
    }
    fit <- sum(res[])							# Sum of squared residuals from true data
    fit.new <- sum(res.new[])						# Sum of squared residuals from ideal data
    test <- step(fit.new - fit)						# Tests whether new data set more extreme
    bpvalue <- mean(test)						# Bayesian p-value
}"
writeLines(jags_model , con = "./Scripts/Fecundity/ZIP.jags" )

#===========================================================================================================================================================
#	DATA IMPORT AND PREPARATION
#===========================================================================================================================================================
### Import data on families
Data <- read.table(file = "./Data/Family.txt", header = TRUE, sep = "\t", dec = ".", na.strings = "NA")
# Average parent body size
Data$Parentsdl <- (Data$Mothersdl + Data$Fathersdl)/2

# Clean and reset factor levels
Data <- Data[!is.na(Data$Mother.inb),]
Data <- Data[Data$Generation != "F1" & Data$Generation != "F2",]
Data$Line <- factor(Data$Line); Data$Generation <- factor(Data$Generation);

### Bundle data for JAGS
jags.data <- list(C = Data$NbEggs, offset = log(Data$Nb_spawning_days), Line = as.integer(Data$Line), Gener = as.integer(Data$Generation), Father.inb = (Data$Father.inb - mean(Data$Father.inb, na.rm = TRUE)) / sd(Data$Father.inb, na.rm = TRUE), Mother.inb = (Data$Mother.inb - mean(Data$Mother.inb, na.rm = TRUE)) / sd(Data$Mother.inb, na.rm = TRUE), Mother.sdl = (log(Data$Mothersdl) - mean(log(Data$Mothersdl), na.rm = TRUE)) / sd(log(Data$Mothersdl), na.rm = TRUE), Father.sdl = (Data$Fathersdl - mean(Data$Fathersdl, na.rm = TRUE)) / sd(Data$Fathersdl, na.rm = TRUE), Parent.sdl = (Data$Parentsdl - mean(Data$Parentsdl, na.rm = TRUE)) / sd(Data$Parentsdl, na.rm = TRUE), N = length(Data$NbEggs), nline = length(levels(Data$Line)), ngener = length(levels(Data$Generation)));

#===========================================================================================================================================================
#	PARAMETERS TO MONITOR
#===========================================================================================================================================================
parameters = c("sigma.eps", "sigma.alpha", "sigma.beta", "alpha1", "alpha2[2]", "alpha2[3]", "alpha3", "alpha4", "alpha5", "alpha6", "beta1", "beta2[2]", "beta2[3]", "beta3", "beta4", "beta5", "beta6", "bpvalue")

#===========================================================================================================================================================
#	MCMC SETTINGS
#===========================================================================================================================================================
ni <- 45000
nt <- 5
nb <- 40000
nc <- 3

#===========================================================================================================================================================
#	MCMC SETTINGS
#===========================================================================================================================================================
inits <- function(){  
       list(u = rep(0, length(Data$NbEggs)))  
     }

#===========================================================================================================================================================
#	SAMPLING POSTERIOR
#===========================================================================================================================================================
library(jagsUI)
mod <- jags(data = jags.data, parameters.to.save = parameters, model.file = "./Scripts/Fecundity/ZIP.jags", n.chains = nc, n.thin = nt, n.iter = ni, n.burnin = nb, n.adapt = 2000, parallel = TRUE) #
```
