## Supplementary material for "Unidirectional response to bidirectional selection on body size. I. Phenotypic, life history and endocrine response": Multivariate_normal_group_VCOV.R

```
#==========================================================================================================================================================
#			R script for Unidirectional response to bidirectional selection on body size. I. Phenotypic, life history and endocrine response.
#			Eric Edeline July 2020
#
#			MULTIVARIATE NORMAL MODEL IN JAGS -- EFFECT PARAMETERIZATION
#
#			/!\ BEFORE RUNNING:	SET WORKING DIRECTORY TO THE Data_and_code FOLDER ON YOUR OWN MACHINE
#						FOR INSTANCE ON MY MACHINE: setwd("~/Dropbox/Article_Clem/Manuscript/Submission/Ecology_and_Evolution/Data_and_codes")
#==================================================================================================================================================
######## FULL MODEL:
modelstring = "
model
{
  ### Likelihood 
  for(i in 1:nfish){
	CC[i, 1:ngen] ~ dmnorm(mu[i, 1:ngen], Sigma.inv[Line[i], 1:ngen, 1:ngen])
	mu[i, 1:ngen] <- offset[i] + alpha0[Generation[i],1:ngen] + alpha1[1:ngen] + beta0[Line[i],1:ngen] + beta1[Sex[i],1:ngen] + beta2[1:ngen] * Inb[i] + beta3[1:ngen] * Sdl[i]
	}

  ### Priors
    # prior for the variance-covariance matrix
      for (i in 1:nline){
	Sigma.inv[i, 1:ngen, 1:ngen] ~ dwish(R[,], ngen)	
	Sigma[i, 1:ngen, 1:ngen] <- inverse(Sigma.inv[i, 1:ngen, 1:ngen])
      }

  for(i in 1:ngen){
     for(j in 1:ngeneration){		# Generation-level random effect in each gene
	alpha0[j,i] ~ dnorm(0.0, tau.alpha0[i])
	}
     tau.alpha0[i] <- pow(sd.alpha0[i], -2)
     sd.alpha0[i] ~ dunif(0,100)
     alpha1[i] ~ dnorm(0.0, 0.01)	# Intercept: Line A, Sex M, average Inb, average Sdl
     beta0[1,i] <- 0.0
     beta0[2,i] ~ dnorm(0.0, 0.01)	# Line G effect in sex M
     beta0[3,i] ~ dnorm(0.0, 0.01)	# Line P effect in sex M
     beta1[1,i] <- 0.0
     beta1[2,i] ~ dnorm(0.0, 0.01)	# Sex F effect in A line
     beta1[3,i] ~ dnorm(0.0, 0.01)	# Sex I effect in A line
     beta2[i] ~ dnorm(0.0, 0.01)	# Slope of Inb effect
     beta3[i] ~ dnorm(0.0, 0.01)	# Slope of Sdl effect

  }
  # Second slot of prior on sex for missing sex data
  for(i in 1:nsex){
	sexprob[i] ~ dunif(0, 1)	
  }
#### Compute derived quantities
  for(i in 1:nline){
    res.LH[i] <- Sigma[i, 1,1]	# Residual standard deviation for each gene in each line
    res.FSH[i] <- Sigma[i, 2,2]
    res.GH[i] <- Sigma[i, 3,3]
    Corr.LH.FSH[i] <- Sigma[i, 1,2] / sqrt(Sigma[i, 1,1] *  Sigma[i, 2,2])	# Residual correlations between genes in each line
    Corr.LH.GH[i] <- Sigma[i, 1,3] / sqrt(Sigma[i, 1,1] *  Sigma[i, 3,3])
    Corr.FSH.GH[i] <- Sigma[i, 2,3] / sqrt(Sigma[i, 2,2] *  Sigma[i, 3,3])
  }
# Assess model fit using sums-of-squares type of discrepancy
  for(i in 1:nfish){
    sq[i,1:ngen] <- pow(CC[i,1:ngen] - mu[i,1:ngen], 2)			#squared residuals
    # Generate replicate data and compute fit stats for them
    CC.new[i,1:ngen] ~ dmnorm(mu[i,1:ngen], Sigma.inv[Line[i], ,])		#ideal data produced from model parameters
    sq.new[i,1:ngen] <- pow(CC.new[i,1:ngen] - mu[i,1:ngen], 2)		#squared residuals from ideal data
  }
  for(j in 1:ngen){
    fit[j] <- sum(sq[,j])			#sum of squared residuals from true data
    fit.new[j] <- sum(sq.new[,j])		#sum of squared residuals from ideal data
    test[j] <- step(fit.new[j] - fit[j])		#test whether new data set more extreme
    bpvalue[j] <- mean(test[j])		#Bayesian p-value
  }
}"
writeLines(modelstring,con="./Scripts/Hormones/Multivariate_normal_group_VCOV.jags")

######## PARAMETERS MONITORED
parameters <- c("res.LH", "res.FSH", "res.GH", "sd.alpha0", "alpha1", "beta0", "beta1", "beta2", "beta3", "Corr.LH.FSH", "Corr.LH.GH", "Corr.FSH.GH", "sd.generation", "bpvalue")

######## DATA PACKAGING
Data <- read.table(file = "./Data/Hormones_multivariate_formatted.txt", sep = "\t", header = TRUE, dec = ".", numerals = "no.loss")
# Remove data prior to F3
Data$Gen <- as.numeric(substr(Data$Generation, 2, 2)); Data <- Data[Data$Gen > 2,]; 
# Remove data from F8, which in fact are old F7 fish
Data <- Data[Data$Gen != 8,]
# Remove missing hormone data (partially-observed nodes not allowed in JAGS)
Data <- Data[!is.na(Data$LH),]; Data <- Data[!is.na(Data$FSH),]; Data <- Data[!is.na(Data$GH),]; Data <- Data[!is.na(Data$Act),]; 
# Reset factor levels
Data$Generation <- factor(Data$Generation); Data$Line <- factor(Data$Line);

# !!!!!!!!!!!!!!!!!!!!!!!!!!!!!!!!!!!!!!!!!!!!!
### Choose here the reference level for sex
Data$Sex <- relevel(Data$Sex, ref = "M")

# Bundle data for JAGS
jags.data <- list(CC = cbind(log(Data$LH), log(Data$FSH), log(Data$GH)), offset = log(Data$Act), Sdl = (Data$Sdl - mean(Data$Sdl, na.rm = TRUE)) / sd(Data$Sdl, na.rm = TRUE), Line = as.numeric(Data$Line), Sex = as.numeric(Data$SexD),  Inb = (Data$Inb - mean(Data$Inb, na.rm = TRUE)) / sd(Data$Inb, na.rm = TRUE), Generation = as.numeric(Data$Generation), nfish = dim(Data)[1], ngen = 3, nline = length(levels(Data$Line)), nsex = length(levels(Data$Sex)), ngeneration = length(levels(Data$Generation)), R = matrix(c(3,0,0,0,3,0,0,0,3), nrow = 3, ncol = 3));

######## MCMC SETTINGS
ni <- 30000
nt <- 5
nb <- 25000
nc <- 3
######## SAMPLING
library(jagsUI)
mod <- jags(data = jags.data, parameters.to.save = parameters, model.file = "./Scripts/Hormones/Multivariate_normal_group_VCOV.jags", n.chains = nc, n.thin = nt, n.iter = ni, n.burnin = nb, n.adapt = 2000, parallel = TRUE); #

# If needed
mod <- update(mod, n.iter = 10000)
```
