## Supplementary material for "Unidirectional response to bidirectional selection on body size. I. Phenotypic, life history and endocrine response": 1.Ogives.R

```
#==========================================================================================================================================================
#			R script for Unidirectional response to bidirectional selection on body size. I. Phenotypic, life history and endocrine response.
#			Eric Edeline July 2020
#
#			JAGS MODEL TO COMPUTE MATURITY OGIVES FOR UNMARKED MEDAKA IN THE LABORATORY -- EFFECTS PARAMETERIZATION
#
#			/!\ BEFORE RUNNING:	SET WORKING DIRECTORY TO THE Data_and_code FOLDER ON YOUR OWN MACHINE
#						FOR INSTANCE ON MY MACHINE: setwd("~/Dropbox/Article_Clem/Manuscript/Submission/Ecology_and_Evolution/Data_and_codes")
#==========================================================================================================================================================
######## 1) MODEL
modelstring = "
 model{
##likelihood
for ( i in 1:n){
  y[i] ~ dbern(p[i])
  logit(p[i]) <-  alpha0 + alpha1[Gen[i]] + alpha2[Line[i]] + (beta0 + beta1[Line[i]]) * Sdl[i] + (beta2 + beta3[Line[i]]) * Age[i] + beta4 * Inb[i] 	
}
### priors
# fixed effects alphas, Cauchy priors following Gelman et al. 2008
  alpha0 ~ dt(0.0, 1/(5^2), 1)		# Intercept for A line
  alpha2[1] <- 0			# Constraint for the effect parameterization
  alpha2[2] ~ dt(0.0, 1/(2.5^2), 1)		# Intercept difference for G line
  alpha2[3] ~ dt(0.0, 1/(2.5^2), 1)		# Intercept difference for P line
# fixed effects betas
  beta0 ~ dt(0.0, 1/(5^2), 1)		# Slope of Sdl effect in the A line
  beta1[1] <- 0				# Constraint
  beta1[2] ~ dt(0.0, 1/(2.5^2), 1)		# Slope difference of Sdl effect in the G line
  beta1[3] ~ dt(0.0, 1/(2.5^2), 1)		# Slope difference of Sdl effect in the P line
#
  beta2 ~ dt(0.0, 1/(5^2), 1)		# Slope of Age effect in the A line
  beta3[1] <- 0				# Constraint
  beta3[2] ~ dt(0.0, 1/(2.5^2), 1)		# Slope difference of Age effect in the G line
  beta3[3] ~ dt(0.0, 1/(2.5^2), 1)		# Slope difference of Age effect in the P line
  beta4 ~ dt(0.0, 1/(5^2), 1)			# Slope of inbreeding effect, same for all lines
# random effect alpha
for(i in 1:ngen){
  alpha1[i] ~ dnorm(0, tau1)		# Random intercept by generation
}
sd1 ~ dunif(0, 100)			
tau1 <- pow(sd1, -2)
### fit assessment
for (i in 1:n){
  resid[i] <- (y[i]-p[i])					#raw residuals
  y.new[i] ~ dbern(p[i]);					#create replicate dataset ideal because created from model parameters
  resid.new[i] <- (y.new[i]-p[i])
}
#discrepancy measures
fit <- sum(resid[]);
fit.new <- sum(resid.new[]);
test <- step(fit.new - fit)		#test whether new data set more extreme
bpvalue <- mean(test)			#Bayesian p-value
}"
writeLines(modelstring,con="./Scripts/PMRN/Ogive_model.jags")

######## PARAMETERS MONITORED
parameters <- c("alpha0", "alpha2", "beta0", "beta1", "beta2", "beta3", "beta4", "sd1", "bpvalue")

######## DATA IMPORT AND CONDITIONING
Data <- read.table(file = "./Data/sdl-filtered.txt", header = TRUE, sep = "\t")
## Compute individual age from mean family birthdate
source("./R_Functions/AgeCorrect.R");
Data$Age <- Age(Data)
## Compute inbreeding coefficient for each individual
source("./R_Functions/Inbreeding.R");
Data$Inb <- Inbreeding(Data, method = "genlib")
## Remove first 2 generations plus first two phenotyping (no mature fish at P0 and P1), and reset factor levels
Data <- Data[Data$Generation != "F1" & Data$Generation != "F2",]; 
Data <- Data[Data$Phenotyping != "P0" & Data$Phenotyping != "P1",]; 
Data$Generation <- factor(Data$Generation); Data$Line <- factor(Data$Line); Data$Phenotyping <- factor(Data$Phenotyping);
# Clean
Data <- Data[!is.na(Data$Sdl),]
Data <- Data[!is.na(Data$Age),]
# Create maturity variable
Data$Mat <- ifelse(Data$Sex == "I", 0, 1);

######## DATA PACKAGING
jags.data <- list(y = Data$Mat, Line = as.integer(Data$Line), Sdl = (Data$Sdl - mean(Data$Sdl)) / (2*sd(Data$Sdl)), Age = (Data$Age - mean(Data$Age)) / (2*sd(Data$Age)), Inb = (Data$Inb - mean(Data$Inb)) / (2*sd(Data$Inb)), Gen = as.integer(Data$Generation), Fam = as.integer(Data$Family), n = length(Data$Mat), nline = 3, ngen = length(levels(Data$Generation)), nfam = length(levels(Data$Family)));

######## MCMC SETTINGS
ni <- 10000
nt <- 5
nb <- 5000
nc <- 3

######## SAMPLING
library(jagsUI)
mod <- jags(data = jags.data, parameters.to.save = parameters, model.file = "./Scripts/PMRN/Ogive_model.jags", n.chains = nc, n.thin = nt, n.iter = ni, n.burnin = nb, n.adapt = 2000, parallel = TRUE);
# If needed
mod <- update(mod, n.iter = 10000)
####### SAVE SAMPLES (removing constraints and unnecessary nodes)
write.table(data.frame(mod$sims.list)[, -c(2, 6, 10, 15:16)], file ="./Scripts/PMRN/Ogive.samples.txt", sep = "\t")
```
