## Supplementary material for "Unidirectional response to bidirectional selection on body size. I. Phenotypic, life history and endocrine response": 2.PMRN.R

```
#==========================================================================================================================================================
#			R script for Unidirectional response to bidirectional selection on body size. I. Phenotypic, life history and endocrine response.
#			Eric Edeline July 2020
#
#			CREATE SYNTHETIC GOWTH CURVES FOR VISUALIZING PMRNs
#
#			/!\ BEFORE RUNNING:	SET WORKING DIRECTORY TO THE Data_and_code FOLDER ON YOUR OWN MACHINE
#						FOR INSTANCE ON MY MACHINE: setwd("~/Dropbox/Article_Clem/Manuscript/Submission/Ecology_and_Evolution/Data_and_codes")
#==========================================================================================================================================================
######## DATA IMPORT AND CONDITIONING
Data <- read.table(file = "./Data/sdl-filtered.txt", header = TRUE, sep = "\t")
## Compute individual age from mean family birthdate
source("./R_Functions/AgeCorrect.R");
Data$Age <- Age(Data)
## Remove first 2 generations and reset factor levels
Data <- Data[Data$Generation != "F1" & Data$Generation != "F2",];
Data$Generation <- factor(Data$Generation); Data$Line <- factor(Data$Line); Data$Phenotyping <- factor(Data$Phenotyping);
# Clean
Data <- Data[!is.na(Data$Sdl),]
Data <- Data[!is.na(Data$Age),]
# Create maturity variable
Data$Mat <- ifelse(Data$Sex == "I", 0, 1);
# Scale length and age to work on the same scale (units) as the ogive model
Data$Sdl <- (Data$Sdl - mean(Data$Sdl)) / (2*sd(Data$Sdl[Data$Phenotyping != "P0" & Data$Phenotyping != "P1"])); 
Data$Age <- (Data$Age - mean(Data$Age)) / (2*sd(Data$Age[Data$Phenotyping != "P0" & Data$Phenotyping != "P1"]));

# Approximate growth curves for slow, medium and fast growers using cubic splines
library(mgcv)
g.med <- gam(Sdl ~ s(Age, k = 4), data = Data)
g.fast <- gam(Sdl ~ s(Age, k = 4), data = rbind(Data[Data$Phenotyping == "P0",], Data[Data$Phenotyping == "P1" & Data$Sdl > quantile(Data$Sdl[Data$Phenotyping == "P1"], probs = 0.75),], Data[Data$Phenotyping == "P1c" & Data$Sdl > quantile(Data$Sdl[Data$Phenotyping == "P1"], probs = 0.75),], Data[Data$Phenotyping == "P2" & Data$Sdl > quantile(Data$Sdl[Data$Phenotyping == "P2"], probs = 0.75),], Data[Data$Phenotyping == "P3" & Data$Sdl > quantile(Data$Sdl[Data$Phenotyping == "P3"], probs = 0.75),]))
g.slow <- gam(Sdl ~ s(Age, k = 4), data = rbind(Data[Data$Phenotyping == "P0",], Data[Data$Phenotyping == "P1" & Data$Sdl < quantile(Data$Sdl[Data$Phenotyping == "P1"], probs = 0.25),], Data[Data$Phenotyping == "P1" & Data$Sdl < quantile(Data$Sdl[Data$Phenotyping == "P1c"], probs = 0.25),], Data[Data$Phenotyping == "P2" & Data$Sdl < quantile(Data$Sdl[Data$Phenotyping == "P2"], probs = 0.25),], Data[Data$Phenotyping == "P3" & Data$Sdl < quantile(Data$Sdl[Data$Phenotyping == "P3"], probs = 0.25),]))

## Visualize the growth curves
plot(Data$Sdl ~ Data$Age, col = "grey70")
lines(predict(g.med, newdata = data.frame(Age = seq(min(Data$Age), max(Data$Age), length.out = 200))) ~ seq(min(Data$Age), max(Data$Age), length.out = 200))
lines(predict(g.slow, newdata = data.frame(Age = seq(min(Data$Age), max(Data$Age), length.out = 200))) ~ seq(min(Data$Age), max(Data$Age), length.out = 200), col = "blue")
lines(predict(g.fast, newdata = data.frame(Age = seq(min(Data$Age), max(Data$Age), length.out = 200))) ~ seq(min(Data$Age), max(Data$Age), length.out = 200), col = "red")

## Build the 3 growth curves and bundle in a single matrix: column 1 = slow growth, column 2 = medium growth, column 3 = fast growth
nage <- 200	# Choose here number of age increments
Age <- seq(min(Data$Age), max(Data$Age), length.out = nage)	
# Save for a plot later
write.table(Age, file = "./Scripts/PMRN/Age.txt", sep ="\t", row.names = FALSE, col.names = FALSE)
Sdl <- cbind(predict(g.slow, newdata = data.frame(Age = Age)), predict(g.med, newdata = data.frame(Age = Age)), predict(g.fast, newdata = data.frame(Age = Age)))
# Save for a later plot
write.table(Sdl, file = "./Scripts/PMRN/Sdl.txt", sep ="\t", row.names = FALSE, col.names = FALSE)

#==========================================================================================================================================================
#	 IMPORT MCMC SAMPLES FOR OGIVES !!! MODEL FITTED ON SCALED SDL AND AGE !!!
#
#			/!\ BEFORE RUNNING:	SET WORKING DIRECTORY TO THE Data_and_code FOLDER ON YOUR OWN MACHINE
#						FOR INSTANCE ON MY MACHINE: setwd("~/Dropbox/Article_Clem/Manuscript/Submission/Ecology_and_Evolution/Data_and_codes")
#==========================================================================================================================================================
Samps <- read.table(file = "./Scripts/PMRN/Ogive.samples.txt", header = TRUE, sep = "\t")
# 1 = A line, 2 = G line, 3 = P line, 
# alpha0 = Model intercept, A line
# alpha2.2 = Intercept difference for G line,
# alpha2.3 = Intercept difference for P line
# beta0 = Slope of Sdl effect in the A line
# beta1.2 = Slope difference of Sdl effect in the G line
# beta1.3 = Slope difference of Sdl effect in the P line
# beta2 = Slope of Age effect in the A line
# beta3.2 = Slope difference of Age effect in the G line
# beta3.3 = Slope difference of Age effect in the P line
#==========================================================================================================================================================
#	 CALCULATION OF PMRN AND LENGTH AT 50% MATURATION PROBABILITY
#==========================================================================================================================================================
## Utility functions
invlogit <- function(x) {1/(1+exp(-x))}
logit <- function(x) {log(x/(1-x))}

###### COMPUTE PMRNs
#### Storage matrices for 50% maturation probability
lp50.C <- matrix(NA, nrow = dim(Samps)[1], ncol = dim(Sdl)[2]); # Storage matrix for length at 50% maturation probability in Control line
ap50.C <- matrix(NA, nrow = dim(Samps)[1], ncol = dim(Sdl)[2]); # Storage matrix for age at 50% maturation probability in Control line
lp50.L <- matrix(NA, nrow = dim(Samps)[1], ncol = dim(Sdl)[2]); # Storage matrix for length at 50% maturation probability in Low line
ap50.L <- matrix(NA, nrow = dim(Samps)[1], ncol = dim(Sdl)[2]); # Storage matrix for age at 50% maturation probability in Low line
lp50.H <- matrix(NA, nrow = dim(Samps)[1], ncol = dim(Sdl)[2]); # Storage matrix for length at 50% maturation probability in High line
ap50.H <- matrix(NA, nrow = dim(Samps)[1], ncol = dim(Sdl)[2]); # Storage matrix for age at 50% maturation probability in High line

for(i in 1:dim(Samps)[1]){		# Loop over MCMC samples for ogive parameters
    for(j in 1:dim(Sdl)[2]){		# Loop over the 3 growth curves
	###### CONTROL LINE
        # Maturity prob
	P.maturity.C <- invlogit(Samps$alpha0[i] + Samps$beta0[i] * matrix(Sdl[,j]) + Samps$beta2[i] * Age)		
        # Maturation prob from Eq. 2 page 663 in Barot et al. 2004. P.maturation.C can start being computed from 2nd age, hence use P.maturity.C[-1]
  	P.maturation.C <- (P.maturity.C[-1] - P.maturity.C[-length(P.maturity.C)]) / (1 - P.maturity.C[-length(P.maturity.C)])	
	# Length at 50% maturation probability from Eq. 5 page 6 in Harney at al 2013
  	lp50.C[i,j] <- Sdl[-1, j][min(which(1-cumprod(1-P.maturation.C) >= 0.5))]	# Sdl[-1,] to match the size of maturation vector	
	# Age at 50% maturation probability from Eq. 5 page 6 in Harney at al 2013
  	ap50.C[i,j] <- Age[-1][min(which(1-cumprod(1-P.maturation.C) >= 0.5))]		# Age[-1,] to match the size of maturation vector

	###### LOW LINE
	P.maturity.L <- invlogit(Samps$alpha0[i] + Samps$alpha2.3[i] + (Samps$beta0[i] + Samps$beta1.3[i]) * matrix(Sdl[,j]) + (Samps$beta2[i] + Samps$beta3.3[i]) * Age)
  	P.maturation.L <- (P.maturity.L[-1] - P.maturity.L[-length(P.maturity.L)]) / (1 - P.maturity.L[-length(P.maturity.L)])
  	lp50.L[i,j] <- Sdl[-1, j][min(which(1-cumprod(1-P.maturation.L) >= 0.5))]	
  	ap50.L[i,j] <- Age[-1][min(which(1-cumprod(1-P.maturation.L) >= 0.5))]
		
	##### HIGH LINE
	P.maturity.H <- invlogit(Samps$alpha0[i] +Samps$alpha2.2[i] + (Samps$beta0[i] + Samps$beta1.2[i]) * matrix(Sdl[,j]) + (Samps$beta2[i] + Samps$beta3.2[i])* Age)
  	P.maturation.H <- (P.maturity.H[-1] - P.maturity.H[-length(P.maturity.H)]) / (1 - P.maturity.H[-length(P.maturity.H)])
  	lp50.H[i,j] <- Sdl[-1, j][min(which(1-cumprod(1-P.maturation.H) >= 0.5))]	
  	ap50.H[i,j] <- Age[-1][min(which(1-cumprod(1-P.maturation.H) >= 0.5))]		
  }
}
## Warnings are produced if 50% probability is not reached
#### SAVE THE PMRN DATA
write.table(lp50.C, file = "./Scripts/PMRN/Lp50_C.txt", sep = "\t") 
write.table(ap50.C, file = "./Scripts/PMRN/Ap50_C.txt", sep = "\t") 
write.table(lp50.L, file = "./Scripts/PMRN/Lp50_L.txt", sep = "\t") 
write.table(ap50.L, file = "./Scripts/PMRN/Ap50_L.txt", sep = "\t") 
write.table(lp50.H, file = "./Scripts/PMRN/Lp50_H.txt", sep = "\t") 
write.table(ap50.H, file = "./Scripts/PMRN/Ap50_H.txt", sep = "\t") 
```
