## Supplementary material for "Unidirectional response to bidirectional selection on body size. I. Phenotypic, life history and endocrine response": 3.Plot_PMRN.R

```
#==========================================================================================================================================================
#			R script for Unidirectional response to bidirectional selection on body size. I. Phenotypic, life history and endocrine response.
#			Eric Edeline July 2020
#
#	 		PLOT THE PMRNs
#
#			/!\ BEFORE RUNNING:	SET WORKING DIRECTORY TO THE Data_and_code FOLDER ON YOUR OWN MACHINE
#						FOR INSTANCE ON MY MACHINE: setwd("~/Dropbox/Article_Clem/Manuscript/Submission/Ecology_and_Evolution/Data_and_codes")
#==========================================================================================================================================================
######## DATA IMPORT AND CONDITIONING
Data <- read.table(file = "./Data/sdl-filtered.txt", header = TRUE, sep = "\t")
## Compute individual age from mean family birthdate
source("./R_Functions/AgeCorrect.R");
Data$Age <- Age(Data)
## Remove first 2 generations and reset factor levels
Data <- Data[Data$Generation != "F1" & Data$Generation != "F2",]; 
Data$Generation <- factor(Data$Generation); Data$Line <- factor(Data$Line); Data$Phenotyping <- factor(Data$Phenotyping);
# Clean
Data <- Data[!is.na(Data$Sdl),]
Data <- Data[!is.na(Data$Age),]

######## IMPORT PMRN SAMPLES
Lp50_C <- read.table(file = "./Scripts/PMRN/Lp50_C.txt", sep = "\t")
Ap50_C <- read.table(file = "./Scripts/PMRN/Ap50_C.txt", sep = "\t")
Lp50_L <- read.table(file = "./Scripts/PMRN/Lp50_L.txt", sep = "\t")
Ap50_L <- read.table(file = "./Scripts/PMRN/Ap50_L.txt", sep = "\t")
Lp50_H <- read.table(file = "./Scripts/PMRN/Lp50_H.txt", sep = "\t")
Ap50_H <- read.table(file = "./Scripts/PMRN/Ap50_H.txt", sep = "\t")

####### BACK-SCALING FUNCTIONS
Sdl.sc <- function(x) {x * 2 *sd(Data$Sdl[Data$Phenotyping != "P0" & Data$Phenotyping != "P1"], na.rm = TRUE) + mean(Data$Sdl, na.rm = TRUE)}
Age.sc <- function(x) {x * 2 * sd(Data$Age[Data$Phenotyping != "P0" & Data$Phenotyping != "P1"], na.rm = TRUE) + mean(Data$Age, na.rm = TRUE)}
####### BACK-SCALE DATA
Sdl <- Sdl.sc(read.table(file = "./Scripts/PMRN/Sdl.txt", sep ="\t", header = FALSE))
Age <- Age.sc(read.table(file = "./Scripts/PMRN/Age.txt", sep ="\t", header = FALSE))
Lp50.C <- Sdl.sc(Lp50_C)
Ap50.C <- Age.sc(Ap50_C)
Lp50.L <- Sdl.sc(Lp50_L)
Ap50.L <- Age.sc(Ap50_L)
Lp50.H <- Sdl.sc(Lp50_H)
Ap50.H <- Age.sc(Ap50_H)

######## 50% MATURATION PROBABILITY PLOT

pdf(file = "./Scripts/PMRN/PMRN_F3_to_F7.pdf")
par(omi = c(0,0,0,0), mai = c(.9,.9,0,0))
plot(Data$Sdl ~ Data$Age, xlab = "Age (days)", ylab = "Standard length (mm)", main = "", cex.lab = 1.3, axes = FALSE, type = "n")
# Line-specific raw data
points(Data$Sdl[Data$Line == "A"] ~ Data$Age[Data$Line == "A"], col = "grey80", pch = 1);
points(Data$Sdl[Data$Line == "G"] ~ Data$Age[Data$Line == "G"], col = "grey80", pch = 2);
points(Data$Sdl[Data$Line == "P"] ~ Data$Age[Data$Line == "P"], col = "grey80", pch = 6);
# Synthetic growth curves
lines(Sdl$V3 ~ Age$V1, lty = 2); lines(Sdl$V2 ~ Age$V1, lty = 2); lines(Sdl$V1 ~ Age$V1, lty = 2);

# 50 % maturation segments
segments(x0 = median(Ap50.C[,3], na.rm = TRUE), y0 = median(Lp50.C[,3], na.rm = TRUE), x1 = median(Ap50.C[,1], na.rm = TRUE), y1 = median(Lp50.C[,1], na.rm = TRUE), lwd = 1)
segments(x0 = median(Ap50.L[,3], na.rm = TRUE), y0 = median(Lp50.L[,3], na.rm = TRUE), x1 = median(Ap50.L[,1], na.rm = TRUE), y1 = median(Lp50.L[,1], na.rm = TRUE), lwd = 1, col = "blue")
segments(x0 = median(Ap50.H[,3], na.rm = TRUE), y0 = median(Lp50.H[,3], na.rm = TRUE), x1 = median(Ap50.H[,1], na.rm = TRUE), y1 = median(Lp50.H[,1], na.rm = TRUE), lwd = 1, col = "red")
# Length errors
segments(x0 = apply(Ap50.C, 2, median, na.rm = TRUE), y0 = apply(Lp50.C, 2, quantile, probs = 0.025, na.rm = TRUE), x1 = apply(Ap50.C, 2, median, na.rm = TRUE), y1 = apply(Lp50.C, 2, quantile, probs = 0.975, na.rm = TRUE), lwd = 1)
segments(x0 = apply(Ap50.L, 2, median, na.rm = TRUE), y0 = apply(Lp50.L, 2, quantile, probs = 0.025, na.rm = TRUE), x1 = apply(Ap50.L, 2, median, na.rm = TRUE), y1 = apply(Lp50.L, 2, quantile, probs = 0.975, na.rm = TRUE), col = "blue", lwd = 1)
segments(x0 = apply(Ap50.H, 2, median, na.rm = TRUE), y0 = apply(Lp50.H, 2, quantile, probs = 0.025, na.rm = TRUE), x1 = apply(Ap50.H, 2, median, na.rm = TRUE), y1 = apply(Lp50.H, 2, quantile, probs = 0.975, na.rm = TRUE), col = "red", lwd = 1)
# Age errors
#segments(x0 = apply(Ap50.C, 2, quantile, probs = 0.025, na.rm = TRUE), y0 = apply(Lp50.C, 2, median, na.rm = TRUE), x1 = apply(Ap50.C, 2, quantile, probs = 0.975, na.rm = TRUE), y1 = apply(Lp50.C, 2, median, na.rm = TRUE), lwd = 2)
#segments(x0 = apply(Ap50.L, 2, quantile, probs = 0.025, na.rm = TRUE), y0 = apply(Lp50.L, 2, median, na.rm = TRUE), x1 = apply(Ap50.L, 2, quantile, probs = 0.975, na.rm = TRUE), y1 = apply(Lp50.L, 2, median, na.rm = TRUE), lwd = 2, col = "blue")
#segments(x0 = apply(Ap50.H, 2, quantile, probs = 0.025, na.rm = TRUE), y0 = apply(Lp50.H, 2, median, na.rm = TRUE), x1 = apply(Ap50.H, 2, quantile, probs = 0.975, na.rm = TRUE), y1 = apply(Lp50.H, 2, median, na.rm = TRUE), lwd = 2, col = "red")
# Points
points(x = apply(Ap50.C, 2, median, na.rm = TRUE), y = apply(Lp50.C, 2, median, na.rm = TRUE))
points(x = apply(Ap50.L, 2, median, na.rm = TRUE), y = apply(Lp50.L, 2, median, na.rm = TRUE), col = "blue", pch = 6)
points(x = apply(Ap50.H, 2, median, na.rm = TRUE), y = apply(Lp50.H, 2, median, na.rm = TRUE), col = "red", pch = 2)
axis(side = 1); axis(side = 2);
dev.off()
```
