## Supplementary material for "Unidirectional response to bidirectional selection on body size. I. Phenotypic, life history and endocrine response": Sdl_0dph.R

```
#==========================================================================================================================================================
#			R script for Unidirectional response to bidirectional selection on body size. I. Phenotypic, life history and endocrine response.
#			Eric Edeline July 2020
#
#			JAGS MODEL FOR SDL AT HATCH -- EFFECT PARAMETRIZATION
#
#			/!\ BEFORE RUNNING:	SET WORKING DIRECTORY TO THE Data_and_code FOLDER ON YOUR OWN MACHINE
#						FOR INSTANCE ON MY MACHINE: setwd("~/Dropbox/Article_Clem/Manuscript/Submission/Ecology_and_Evolution/Data_and_codes")
#==========================================================================================================================================================
jags_model = "
  model {
# Likelihood
    for (i in 1:N) {
      y[i] ~ dnorm(mu[i], 1/sigma^2)
      mu[i] <- alpha0[Gener[i]] + beta0 + beta1[Line[i]] + beta2 * Pa.Sdl[i] + beta3 * Inb[i]
    }
# Priors:
  sigma ~ dunif(0, 10)
  for(i in 1:ngener){
    alpha0[i] ~ dnorm(0, 1/sigma.alpha0^2)  
  }
  sigma.alpha0 ~ dunif(0, 10)
  beta0 ~ dnorm(0, 0.001)		# Intercept = line A, at an average Mo.Sdl and inbreeding
  beta1[1] <- 0				# Constraint for the effect parametrization
  beta1[2] ~ dnorm(0 , 0.001)  		# Effect of G line
  beta1[3] ~ dnorm(0 , 0.001)		# Effect of P line
  beta2 ~ dnorm(0, 0.001)		# Slope of Pa.Sdl effect
  beta3 ~ dnorm(0, 0.001)		# Slope of inbreeding effect
# Assess model fit using sums-of-squares type of discrepancy
  for(i in 1:N){
    sq[i] <- pow(y[i] - mu[i], 2)			#squared residuals
    # Generate replicate data and compute fit stats for them
    y.new[i] ~ dnorm(mu[i], 1/sigma^2)			#ideal data produced from model parameters
    sq.new[i] <- pow(y.new[i] - mu[i], 2)	#squared residuals from ideal data
    }
    fit <- sum(sq[])			#sum of squared residuals from true data
    fit.new <- sum(sq.new[])		#sum of squared residuals from ideal data
    test <- step(fit.new - fit)		#test whether new data set more extreme
    bpvalue <- mean(test)		#Bayesian p-value
    c.hat <- fit/(fit.new + 0.00001)
} "
writeLines(jags_model , con = "./Scripts/Sdl_at_0_dph/Normal_Sdl_0dph.jags" )

#######################################################################################################################################
#	PARAMETERS TO MONITOR
#######################################################################################################################################
parameters = c("sigma", "sigma.alpha0", "beta0", "beta1", "beta2", "beta3", "bpvalue", "c.hat")

#######################################################################################################################################
#	DATA
#######################################################################################################################################
Data <- read.table(file = "./Data/sdl-filtered.txt", header = TRUE, sep = "\t")
## Compute inbreeding coefficient for each individual
source("./R_Functions/Inbreeding.R");
Data$Inb <- Inbreeding(Data, method = "genlib")
## Add mother and father Sdl data
Data$Mo.Sdl <- Data$Sdl[match(Data$Mother, Data$ID)]
Data$Fa.Sdl <- Data$Sdl[match(Data$Father, Data$ID)]
Data$Pa.Sdl <- (Data$Mo.Sdl + Data$Fa.Sdl)/2

## Remove first 2 generations and keep only first phenotyping
Data <- Data[Data$Generation != "F1" & Data$Generation != "F2",]; 
Data <- Data[Data$Phenotyping == "P0",]; 
Data$Generation <- factor(Data$Generation); Data$Line <- factor(Data$Line); Data$Phenotyping <- factor(Data$Phenotyping);
# Clean
Data <- Data[!is.na(Data$Sdl),]
## Set reference levels
Data$Line <- relevel(Data$Line, ref = "A") 	# choose here the reference level for sex
## Bundle data
jags.data <- list(y = Data$Sdl, N = length(Data$Sdl), Line = as.integer(Data$Line), Mo.Sdl = (Data$Mo.Sdl - mean(Data$Mo.Sdl)) / sd(Data$Mo.Sdl), Pa.Sdl = (Data$Pa.Sdl - mean(Data$Pa.Sdl)) / sd(Data$Pa.Sdl), Inb = (Data$Inb - mean(Data$Inb)) / sd(Data$Inb), Gener = as.integer(Data$Generation), ngener = length(levels(Data$Generation)))

#######################################################################################################################################
#	MCMC SETTINGS
#######################################################################################################################################
ni <- 6000
nt <- 1
nb <- 5000
nc <- 3

#######################################################################################################################################
#	SAMPLING POSTERIOR
#######################################################################################################################################
library(jagsUI)
mod <- jags(data = jags.data, parameters.to.save = parameters, model.file = "./Scripts/Sdl_at_0_dph/Normal_Sdl_0dph.jags", n.chains = nc, n.thin = nt, n.iter = ni, n.burnin = nb, n.adapt = 2000, parallel = TRUE) #
```
