## Supplementary material for "Unidirectional response to bidirectional selection on body size. I. Phenotypic, life history and endocrine response": Sdl_75dph.R

```
#==========================================================================================================================================================
#			R script for Unidirectional response to bidirectional selection on body size. I. Phenotypic, life history and endocrine response.
#			Eric Edeline July 2020
#
#			JAGS MODEL FOR SDL AT P3 -- EFFECT PARAMETRIZATION
#
#			/!\ BEFORE RUNNING:	SET WORKING DIRECTORY TO THE Data_and_code FOLDER ON YOUR OWN MACHINE
#						FOR INSTANCE ON MY MACHINE: setwd("~/Dropbox/Article_Clem/Manuscript/Submission/Ecology_and_Evolution/Data_and_codes")
#==========================================================================================================================================================
jags_model = "
  model {
# Likelihood
  for (i in 1:N) {
    y[i] ~ dnorm(mu[i], 1/sigma^2)
    mu[i] <- alpha0 + alpha1[Gener[i]] + beta0[Line[i]] + beta1[Sex[i]] + beta2 * Age[i] + beta3 * Inb[i]
  }
  sigma ~ dunif(0,10)
# Priors:
  alpha0 ~ dnorm(0, 0.001)		# Intercept = line A, sex F at an average age and inbreeding
  for(i in 1:ngener){
    # Random generation effect
    alpha1[i] ~ dnorm(0, 1/sigma.alpha1^2)  
  }
  sigma.alpha1 ~ dunif(0, 10)
  # Fixed effects
  beta0[1] <- 0				# Constraint for the effect parametrization
  beta0[2] ~ dnorm(0 , 0.001)  		# Effect of G line
  beta0[3] ~ dnorm(0 , 0.001)		# Effect of P line
  beta1[1] <- 0				# Constraint for the effect parameterization
  beta1[2] ~ dnorm(0 , 0.001) 		# Effect of sex I 
  beta1[3] ~ dnorm(0 , 0.001)  		# Effect of sex M
  beta2 ~ dnorm(0, 0.001)		# Slope of age effect
  beta3 ~ dnorm(0, 0.001)		# Slope of inbreeding effect
# Assess model fit using sums-of-squares type of discrepancy
  for(i in 1:N){
    sq[i] <- pow(y[i] - mu[i], 2)			#squared residuals
    # Generate replicate data and compute fit stats for them
    y.new[i] ~ dnorm(mu[i], 1/sigma^2)			#ideal data produced from model parameters
    sq.new[i] <- pow(y.new[i] - mu[i], 2)	#squared residuals from ideal data
    }
    fit <- sum(sq[])			#sum of squared residuals from true data
    fit.new <- sum(sq.new[])		#sum of squared residuals from ideal data
    test <- step(fit.new - fit)		#test whether new data set more extreme
    bpvalue <- mean(test)		#Bayesian p-value
} "
writeLines( jags_model , con = "./Scripts/Sdl_at_75_dph/Normal_Sdl.jags")

#######################################################################################################################################
#	PARAMETERS TO MONITOR
#######################################################################################################################################
parameters = c("sigma", "sigma.alpha1", "alpha0", "beta0", "beta1", "beta2", "beta3", "bpvalue")

#######################################################################################################################################
#	DATA
#######################################################################################################################################
Data <- read.table(file = "./Data/sdl-filtered.txt", header = TRUE, sep = "\t")
## Compute individual age from mean family birthdate
source("./R_Functions/AgeCorrect.R");
Data$Age <- Age(Data)
## Compute inbreeding coefficient for each individual
source("./R_Functions/Inbreeding.R");
Data$Inb <- Inbreeding(Data, method = "genlib")

## Remove first 2 generations plus first two phenotyping (no mature fish at P0 and P1), and reset factor levels
Data <- Data[Data$Generation != "F1" & Data$Generation != "F2",]; 
Data <- Data[Data$Phenotyping != "P0" & Data$Phenotyping != "P1",]; 
Data$Generation <- factor(Data$Generation); Data$Line <- factor(Data$Line); Data$Phenotyping <- factor(Data$Phenotyping);
# Clean
Data <- Data[!is.na(Data$Sdl),]
Data <- Data[!is.na(Data$Age),]
## Keep only data for 3rd phenotyping
Data <- Data[Data$Phenotyping == "P3",]
## Set reference levels
Data$Sex <- relevel(Data$Sex, ref = "F") 	# choose here the reference level for sex
Data$Line <- relevel(Data$Line, ref = "A") 	# choose here the reference level for line
## Bundle data
jags.data <- list(y = Data$Sdl, Age = (Data$Age - mean(Data$Age)) / sd(Data$Age), Line = as.integer(Data$Line), Sex = as.integer(Data$Sex), Inb = (Data$Inb - mean(Data$Inb)) / sd(Data$Inb), Gener = as.integer(Data$Generation), N = length(Data$Sdl), nline = length(levels(Data$Line)), nsex= length(levels(Data$Sex)), ngener = length(levels(Data$Generation)))

#######################################################################################################################################
#	MCMC SETTINGS
#######################################################################################################################################
ni <- 6000
nt <- 1
nb <- 5000
nc <- 3

#######################################################################################################################################
#	SAMPLING POSTERIOR
#######################################################################################################################################
library(jagsUI)
mod <- jags(data = jags.data, parameters.to.save = parameters, model.file = "./Scripts/Sdl_at_75_dph/Normal_Sdl.jags", n.chains = nc, n.thin = nt, n.iter = ni, n.burnin = nb, n.adapt = 2000, parallel = TRUE) #
```
