## Supplementary material for "Unidirectional response to bidirectional selection on body size. I. Phenotypic, life history and endocrine response": Survival.R

```
#==========================================================================================================================================================
#			R script for Unidirectional response to bidirectional selection on body size. I. Phenotypic, life history and endocrine response.
#			Eric Edeline July 2020
#
#			JAGS MODEL FOR SURVIVAL PROBABILITY
#
#			/!\ BEFORE RUNNING:	SET WORKING DIRECTORY TO THE Data_and_code FOLDER ON YOUR OWN MACHINE
#						FOR INSTANCE ON MY MACHINE: setwd("~/Dropbox/Article_Clem/Manuscript/Submission/Ecology_and_Evolution/Data_and_codes")
#==========================================================================================================================================================
modelstring = "
model{
##likelihood
  for ( i in 1 : n){
    C[i] ~ dbin(p[i], N[i])
    logit(p[i]) <- beta0 + beta1[Line[i]] + beta2 * Mother.inb[i] + beta3 * Father.inb[i] + eps[i]
    eps[i] ~ dnorm(0, tau.eps)	# Prior for overdispersion parameter
  }
## prior for overdispersion parameter
  tau.eps <- pow(sigma.eps, -2)
  sigma.eps ~ dunif(0.0, 10)
  
# Priors for fixed effects
    beta0 ~ dt(0.0, 1/(5^2), 1)	# Model intercept = logit(p) in A line
    beta1[1] <- 0		# Constraint
    beta1[2] ~ dt(0.0, 1/(2.5^2), 1)	# Effect of G line compared to A line
    beta1[3] ~ dt(0.0, 1/(2.5^2), 1)	# Effect of P line compared to A line# Priors for fixed effects on lambda
    beta2 ~ dt(0.0, 1/(2.5^2), 1)	# Slope of Mother.inb effect
    beta3 ~ dt(0.0, 1/(2.55^2), 1)	# Slope of Father.inb effect
##fit assessment
  for (i in 1 : n){
    predicted[i] <- N[i]*p[i]
    resid[i] <- (C[i]-N[i]*p[i])/sqrt(N[i]*p[i]*(1-p[i]) + 0.0001)		#Pearson residuals for the real data
    C.new[i] ~ dbin(p[i], N[i])							#create replicate dataset, ideal because created from model parameters
    resid.new[i] <- (C.new[i]-N[i]*p[i])/sqrt(N[i]*p[i]*(1-p[i]) + 0.0001)	#Pearson residuals for the replicate (ideal) data
    D[i] <- pow(resid[i],2)							#squared Pearson residuals
    D.new[i] <- pow(resid.new[i], 2)
  }
#discrepancy measures
  fit <- sum(resid[])
  fit.new <- sum(resid.new[])
  test <- step(fit.new - fit)					# Tests whether new data set more extreme
  bpvalue <- mean(test)						# Bayesian p-value
}"
writeLines(modelstring,con="./Scripts/Survival/Survival.jags")


#==========================================================================================================================================================
#			PARAMETERS MONITORED
#==========================================================================================================================================================
parameters <- c("sigma.eps", "beta0", "beta1[2]", "beta1[3]", "beta2", "beta3", "bpvalue")

#==========================================================================================================================================================
#			DATA IMPORT
#==========================================================================================================================================================
Data <- read.table(file = "./Data/Family.txt", header = TRUE, sep = "\t", dec = ".")
# NbEggs = number of eggs collected per breeding pair
# Nbfry = number of larvae collected from eggs
# Nb_P0 = number of larvae phenotyped and kept to build a sibling family
# Nb_P1 to P3 = number of fish phenotyped at 15, 60 and 75 dph in each sibling family
# Density_egalisation = number of fish kept in the sibling family at P1
# Sample_for_dissection = number of fish dissected between P1 and P2 for hormonal measurement

# Clean and reset factor levels
Data <- Data[!is.na(Data$Mother.inb),]
Data <- Data[Data$Generation != "F1" & Data$Generation != "F2",]
Data$Line <- factor(Data$Line); Data$Generation <- factor(Data$Generation);

### Bundle data for JAGS FOR EGG-TO-LARVAE SURVIVAL (= hatching rate)
jags.data1 <- list(C = Data$Nbfry, N = Data$NbEggs, Line = as.integer(Data$Line), Gener = as.integer(Data$Generation), Father.inb = (Data$Father.inb - mean(Data$Father.inb, na.rm = TRUE)) / (2*sd(Data$Father.inb, na.rm = TRUE)), Mother.inb = (Data$Mother.inb - mean(Data$Mother.inb, na.rm = TRUE)) / (2*sd(Data$Mother.inb, na.rm = TRUE)), n = length(Data$Nbfry), ngener = length(levels(Data$Generation)));

### Bundle data FOR LARVAE-TO-15DPH SURVIVAL
jags.data2 <- list(C = Data$Nb_P1, N = Data$Nb_P0, Line = as.integer(Data$Line), Gener = as.integer(Data$Generation), Father.inb = (Data$Father.inb - mean(Data$Father.inb, na.rm = TRUE)) / (2*sd(Data$Father.inb, na.rm = TRUE)), Mother.inb = (Data$Mother.inb - mean(Data$Mother.inb, na.rm = TRUE)) / (2*sd(Data$Mother.inb, na.rm = TRUE)), n = length(Data$Nb_P1), ngener = length(levels(Data$Generation)));

### Bundle data FOR 15-TO-60DPH SURVIVAL
# Clean data first
Data <- Data[!is.na(Data$Nb_P2),]
Data <- Data[Data$Nb_P2 <= Data$Density_egalisation - Data$Sample_for_dissection,]
jags.data3 <- list(C = Data$Nb_P2, N = Data$Density_egalisation - Data$Sample_for_dissection, Line = as.integer(Data$Line), Gener = as.integer(Data$Generation), Father.inb = (Data$Father.inb - mean(Data$Father.inb, na.rm = TRUE)) / (2*sd(Data$Father.inb, na.rm = TRUE)), Mother.inb = (Data$Mother.inb - mean(Data$Mother.inb, na.rm = TRUE)) / (2*sd(Data$Mother.inb, na.rm = TRUE)), n = length(Data$Nb_P2), ngener = length(levels(Data$Generation)));

### Bundle data FOR 60-TO-75DPH SURVIVAL
# Clean data first
Data <- Data[!is.na(Data$Nb_P2),]
jags.data4 <- list(C = Data$Nb_P3, N = Data$Nb_P2, Line = as.integer(Data$Line), Gener = as.integer(Data$Generation), Father.inb = (Data$Father.inb - mean(Data$Father.inb, na.rm = TRUE)) / (2*sd(Data$Father.inb, na.rm = TRUE)), Mother.inb = (Data$Mother.inb - mean(Data$Mother.inb, na.rm = TRUE)) / (2*sd(Data$Mother.inb, na.rm = TRUE)), n = length(Data$Nb_P3), ngener = length(levels(Data$Generation)));

#===========================================================================================================================================================
#	MCMC SETTINGS
#===========================================================================================================================================================
ni <- 15000
nt <- 5
nb <- 10000
nc <- 3

#===========================================================================================================================================================
#	SAMPLING POSTERIOR
#===========================================================================================================================================================
library(jagsUI)
## !!! Make sure that you choose the right dataset
mod <- jags(data = jags.data4, parameters.to.save = parameters, model.file = "./Scripts/Survival/Survival.jags", n.chains = nc, n.thin = nt, n.iter = ni, n.burnin = nb, n.adapt = 2000, parallel = TRUE)
```
